# Analysis of cardiac differentiation at single cell resolution reveals a requirement of hypertrophic signaling for HOPX transcription

**DOI:** 10.1101/229294

**Authors:** Clayton E Friedman, Quan Nguyen, Samuel W Lukowski, Han Sheng Chiu, Abbigail Helfer, Jason Miklas, Shengbao Suo Suo, Jing-Dong Jackie Han, Pierre Osteil, Guangdun Peng, Naihe Jing, Greg J Baillie, Anne Senabouth, Angelika N Christ, Timothy J Bruxner, Charles E Murry, Emily S Wong, Jun Ding, Yuliang Wang, James Hudson, Hannele Ruohola-Baker, Ziv Bar-Joseph, Patrick P L Tam, Joseph E Powell, Nathan J Palpant

**Affiliations:** Institute for Molecular Bioscience, The University of Queensland, Brisbane, QLD Australia 4072; Centre for Cardiac and Vascular Biology, The University of Queensland, Brisbane, QLD Australia 4072; School of Biomedical Sciences, The University of Queensland, Brisbane, QLD Australia 4072; Key Laboratory of Computational Biology, Chinese Academy of Sciences-Max Planck Partner Institute for Computational Biology, Shanghai Institutes for Biological Sciences, Chinese Academy of Sciences, Shanghai, China; Embryology Unit, Children’s Medical Research Institute, Westmead, NSW 2145 Australia; State Key Laboratory of Cell Biology, CAS Center for Excellence in Molecular Cell Science, Shanghai Institute of Biochemistry and Cell Biology, Chinese Academy of Sciences; University of Chinese Academy of Sciences, 320 Yue Yang Road, Shanghai 200031, China; School of Life Science and Technology, ShanghaiTech University, 100 Haike Road, Shanghai 201210, China; Departments of Pathology, Biochemistry, Bioengineering and Medicine/Cardiology, Institute for Stem Cell and Regenerative Medicine, The University of Washington, Seattle, WA USA; Computational Biology Department, School of Computer Science, Carnegie Mellon University, Pittsburgh, PA 15213, USA; Paul G. Allen School of Computer Science & Engineering, University of Washington, Seattle, WA, 98195; School of Medical Sciences, Sydney Medical School, University of Sydney, NSW 2006 Australia; Garvan-Weizmann Centre for Cellular Genomics, Garvan Institute for Medical Research, Sydney

**Author notes:** co-senior authors and lead contacts (Joseph E Powell, Nathan J Palpant). These authors contributed equally (Clayton E Friedman, Quan Nguyen). Lead contact (N.J.P)).

**Keywords:** human pluripotent stem cells, cardiomyocytes, single cell RNA-seq, heart, development, CRISPR, hypertrophy, lineage tracing, HOPX, *scdiff*

## Abstract

Differentiation into diverse cell lineages requires the orchestration of gene regulatory networks guiding diverse cell fate choices. Utilizing human pluripotent stem cells, we measured expression dynamics of 17,718 genes from 43,168 cells across five time points over a thirty day time-course of *in vitro* cardiac-directed differentiation. Unsupervised clustering and lineage prediction algorithms were used to map fate choices and transcriptional networks underlying cardiac differentiation. We leveraged this resource to identify strategies for controlling *in vitro* differentiation as it occurs *in vivo*. HOPX, a non-DNA binding homeodomain protein essential for heart development *in vivo* was identified as dys-regulated in *in vitro* derived cardiomyocytes. Utilizing genetic gain and loss of function approaches, we dissect the transcriptional complexity of the HOPX locus and identify the requirement of hypertrophic signaling for HOPX transcription in hPSC-derived cardiomyocytes. This work provides a single cell dissection of the transcriptional landscape of cardiac differentiation for broad applications of stem cells in cardiovascular biology.

## Introduction

Studies of cardiac development at single-cell resolution have provided valuable new insights into cell diversity and genetic regulation of cell types revealing mechanisms underlying cardiovascular differentiation and morphogenesis. Single-cell analysis of *in vivo* mouse heart development have revealed chamber-specific and temporal changes in gene expression underlying embryonic heart development from E9.5 to postnatal day 21 establishing anatomical patterns of gene expression in the heart (Li et al., 2016) and new insights into transcriptional programs underlying cardiac maturation (DeLaughter et al., 2016). These studies provide a valuable resource by which to understand transcriptional mechanisms underlying diverse fate choices involved in cardiac development and morphogenesis *in vivo*. Like many single-cell transcriptomic studies, they further highlight the importance of dissecting cell heterogeneity to understand mechanisms underlying the identity and fate of cells in health and disease.

Human pluripotent stem cells are a key model system to study human cardiovascular developmental biology (Murry and Keller, 2008). However, the fidelity by which cardiac directed differentiation *in vitro* recapitulates the transcriptional programs governing the diversity of cell fates generated *in vivo* is not well understood. Analyzing differentiation efficiency has relied extensively on expression signatures from bulk samples consisting of hundreds of thousands of cells which lack the resolution to dissect gene expression and cell subpopulation heterogeneity. Furthermore, identification of rare populations remains challenging. Cardiac progenitor populations, for example, are difficult to identify but important as they constitute cell states underlying decision points in fate diversification (Qyang et al., 2007). Lastly, modelling development and disease requires an accurate, quantifiable analysis of complex decisions underlying the orchestration of heterogeneous cell types responsible for cell phenotypes, from molecular characterization to physiological function.

In this study, we report RNA-sequencing data captured from more than forty thousand single cells navigating stage-specific transitions through *in vitro* cardiac directed differentiation from pluripotency using an established small molecule Wnt modulation protocol (Burridge et al., 2014; Lian et al., 2012). In coordination with a companion paper (Nguyen et al., in review), we utilize the power of this data set to expand our understanding of stem cell directed differentiation as a platform to study cardiovascular development. The overall objective of this study was first to benchmark *in vitro* directed differentiation against *in vivo* development by subpopulation and lineage prediction analysis and second to identify mechanisms for directing *in vitro* differentiation to more accurately model *in vivo* heart development. To this end, using unsupervised clustering analysis of single cell data we identify transcriptionally distinct cell subpopulations transiting cardiac directed differentiation that correlate with mesendoderm fate choices made during gastrulation phases of germ layer specification and progressive developments in cardiovascular differentiation and morphogenesis *in vivo* (Li et al., 2016; Peng et al., 2016). To dissect cell fate choices, we implement *scdiff* (Ding et al., 2018) which is specifically designed for time-course single-cell data, to study transcription factors and their regulatory networks underlying coordinated differentiation of diverse cell subpopulations through differentiation from pluripotency into the cardiac lineage. Since heart development *in vivo* requires instructive cues from exogenous sources like signaling from endoderm and mechanical forces of heart beat and growth, we aimed to leverage single-cell transcriptomic data to identify novel signaling or mechanical strategies for differentiating hPSCs to more accurately pattern cardiac fates. We identify the non-DNA binding homeodomain protein HOPX, a key regulator of heart development (Jain et al., 2015), as dysregulated during differentiation and a potential cause for the immature state of *in vitro* derived cardiomyocytes. We use genetic gain and loss of function hPSCs to show that HOPX is responsive to signals of hypertrophy and is required to drive hypertrophic growth of *in vitro* derived cardiomyocytes. We also dissect the transcriptional complexity of the HOPX locus to show the mechanism for HOPX regulation of hypertrophic growth. Taken together, these data provide a new resource for the community and identify a novel strategy for enhancing derivation of *in vitro* derived cardiomyocytes for applications in cardiovascular biology.

## Results

### Single-cell RNA-sequencing analysis of cardiac directed differentiation

To gain insights into the genetic regulation of cardiovascular development, we performed single-cell transcriptional profiling of human iPSCs navigating from pluripotency through stage-specific transitions in cardiac differentiation (**Figure 1A**). Small molecule Wnt modulation was used as an efficient method to differentiate pluripotent cells toward the cardiac lineage (Burridge et al., 2014; Lian et al., 2012). WTC-CRISPRi hiPSCs (Mandegar et al., 2016) were chosen as the parental cell line for this study. These cells are genetically engineered with an inducible nuclease-dead Cas9 fused to a KRAB repression domain. Transcriptional inhibition by gRNAs targeted to the transcriptional start site is doxycycline-dependent and can be designed to silence genes in an allele-specific manner. The versatility of this line provides a means to use this scRNA-seq data as a reference point for future studies aiming to assess the transcriptional basis of cardiac differentiation at the single-cell level. Cells were verified to have a normal 46 X,Y male karyotype by Giemsa banding analysis before analysis by scRNA-Seq. As with previous time-course genomics studies (Paige et al., 2012; Palpant et al., 2017b), we captured cells at time points corresponding to stage-specific transitions in cell state including pluripotency (day 0), germ layer specification (day 2), and progressing through progenitor (day 5), committed (day 15), and definitive (day 30) cardiac cell states. We harvested a total of 44,020 cells of which 43,168 cells were retained after quality control analysis. In total, we captured expression of 17,718 genes (detected in at least 44 cells and with expression values within the overall expression range of 3 median absolute deviation, as described in our companion paper (Nguyen et al., in review)). We used dimensionality reduction approaches to visualize all 43,168 cells in low-dimensional space (**Figure 1B**), in which cell’s coordinates were estimated so that they preserve the expression similarity (local and global distance in the original multidimensional space) in *t*-SNE plots (left), and the differentiation pseudotime (transition probability between cells) in diffusion plots (right). These data show distinct transcriptomic clustering and distribution of cells undergoing differentiation.

**Figure 1.**
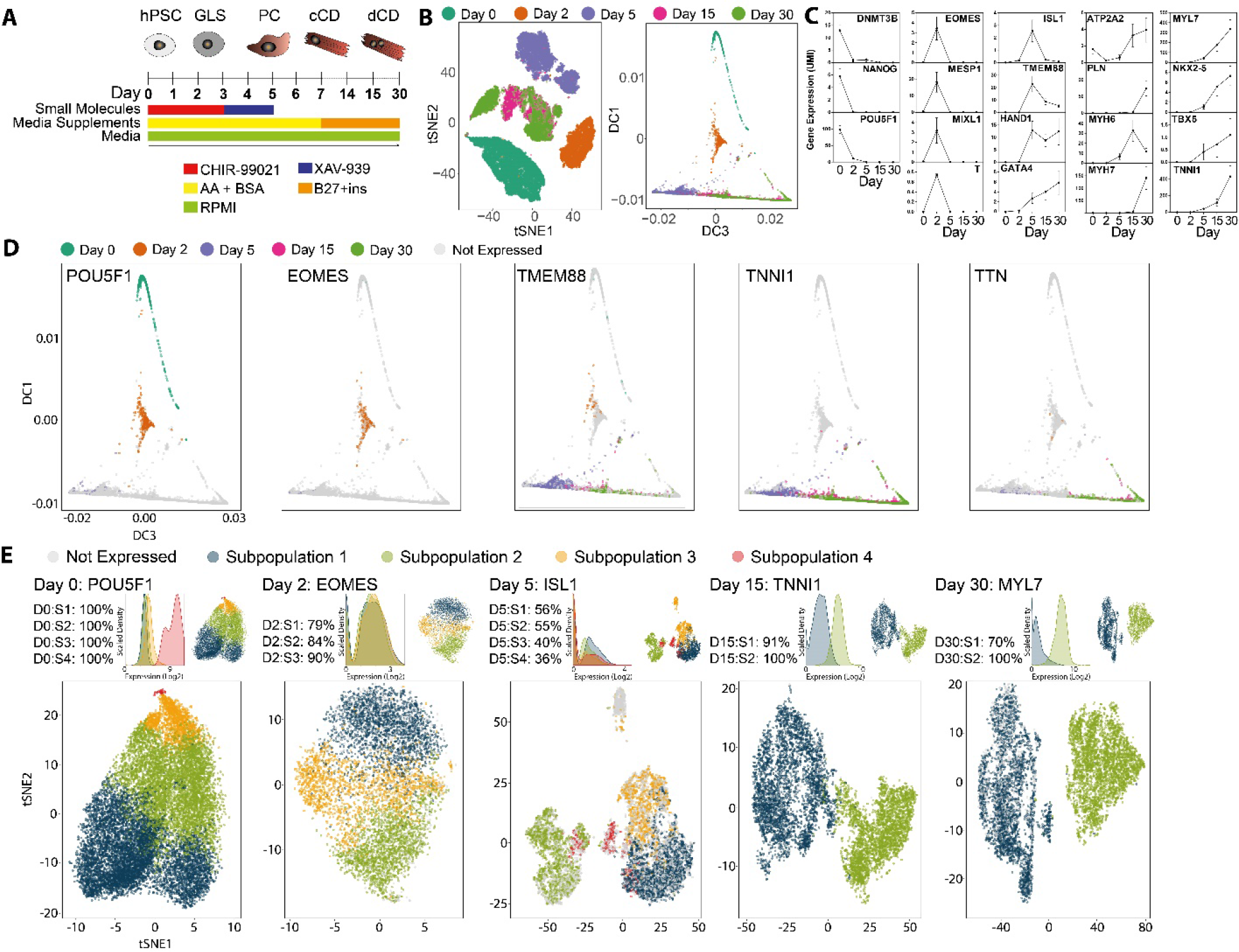
Single-cell analysis of cardiac directed differentiation. **(A)** Schematic of protocol for small molecule directed differentiation from pluripotency into the cardiac lineage. hPSC: human pluripotent stem cell; GLS: germ layer specification; PC: progenitor cell: cCD: committed cardiac derivative; dCD: definitive cardiac derivative. **(B)** 43,168 single-cells transiting cardiac differentiation beginning at pluripotency (day 0) and transitioning through mesoderm (day 2) into progenitor (day 5), committed (day 15), and definitive (day 30) cardiac derivatives. Data are presented using *t*-SNE plot, pseudospacing cells by the nonlinear transformation of similarity in gene expression to preserve the local and global distance of cells in multidimensional space when embedded into two dimensional *t*-SNE space (left), and diffusion plot, pseudospacing cells in a trajectory based on diffusion distance (transition probability) between two cells (right). **(C)** Mean gene expression across all cells at individual time points showing proper temporal expression of stage-specific genes governing differentiation into the cardiac lineage. Shown are pluripotency genes (DNMT3B, POU5F1, NANOG), mes-endoderm genes (EOMES, MIXL1, T, MESP1), and genes governing cardiomyocyte differentiation including signaling regulators (TMEM88), transcription factors (ISL1, HAND1, NKX2-5, TBX5, GATA4), calcium handling genes (ATP2A2, PLN) and sarcomere genes (TNNI1, MYH6, MYH7, MYL7). **(D)** Diffusion plots showing pseudospacing at single-cell resolution for gene expression of stage-specific genes during differentiation based on known genetic regulators of cardiac fate specification including POU5F1 (day 0), EOMES (day 2), TMEM88 (day 5), TNNI1 (day 15), and TTN (day 30). Cells are colored in a binary manner. If the cell expresses the gene it is colored according to the day of isolation (0, 2, 5, 15, or 30). Non-expressing cells are shaded gray. **(E)** Representation of unsupervised clustering analysis (Nguyen et al., in review) using *t*-SNE plots to show single-cell level expression of stage-specific gene expression at each day of differentiation based on known genetic regulators of cardiac fate specification including POU5F1 (day 0), EOMES (day 2), ISL1 (day 5), TNNI1 (day 15), and MYL7 (day 30). If the cell expresses the gene it is colored according to subpopulation 1-4 in which the cell is associated. Non-expressing cells are shaded gray. Above each *t*-SNE plot, the percentage of cells expressing the gene in each subpopulation is shown together with the expression histogram and the reference *t*-SNE plot. Subpopulation coloration across time points does not reflect developmental fate relationship. UMI: unique molecular identifier.

We generated a time-course gene expression profile using a wide range of known cardiac developmental genes by measuring expression among all cells to reveal the temporally-restricted expression dynamics of stage-specific genes reflecting cardiac fate choices (**Figure 1C**). To confirm that the differentiation follows known developmental trajectories, we used dimensionality reduction methods (Coifman et al., 2005; Moignard et al., 2015) (**Figure 1D**) and unsupervised clustering (Clustering at Optimal REsolution (CORE) (Nguyen et al., in review)) (**Figure 1E**) to analyze the expression of known genes governing stage-specific transitions in cardiovascular differentiation at single cell resolution. Overall, these data show that small molecule-mediated cardiac directed differentiation generates developmentally distinct populations of cells displaying expected temporal-specific transcriptional profiles. Our parallel computational genomics study (Nguyen et al., in review) presents a web interface (http://computationalgenomics.com.au/shiny/CardioDifferentiation/), provided as complementary resources for this study.

### Phenotypic diversity and lineage heterogeneity during differentiation

With recent developments in high-resolution transcriptomic mapping of mouse *in vivo* development of the cardiovascular system from the earliest stages of gastrulation (DeLaughter et al., 2016; Li et al., 2016; Peng et al., 2016), we set out to map single-cell heterogeneity of human *in vitro* derived subpopulations to cell types against stages of lineage specification *in vivo*. While cross species comparisons have limitations, this strategy has been used previously for benchmarking *in vitro* differentiation of hPSCs (DeLaughter et al., 2016). To assist in elucidating the molecular identity of each subpopulation, we analyzed high-resolution spatio-temporal gene expression during mouse *in vivo* gastrulation to identify genes that mark known developmental populations and cell types (**Figure S1**). Using previously published approaches, laser microdissection was used to capture germ layer cells of mid-gastrula stage (E7.0) embryos (Peng et al., 2016), with an expanded analysis to include early-(E6.5) and late-gastrulation (E7.5) mouse embryos (unpublished data). High-throughput RNA-sequencing data were compiled into corn plots, with each plot depicting discrete spatial-temporal patterns of gene expression corresponding to individually sequenced sections. To determine phenotypic identities based on gene expression networks governing each human *in vitro-derived* subpopulation during differentiation, we visualized the spatio-temporal patterns of gene expression in the gastrulating mouse embryo including: EOMES (pan-mesendoderm), MESP1 and MIXL1 (mesoderm), SOX17 and FOXA2 (endoderm), and NKX2-5 (cardiac lineage transcription factor) (**Figure 2A**, **Figure S2A-D**). These *in vivo* expression dynamics of mouse gastrulation established spatiotemporal reference points for identifying *in vitro* subpopulations.

**Figure 2.**
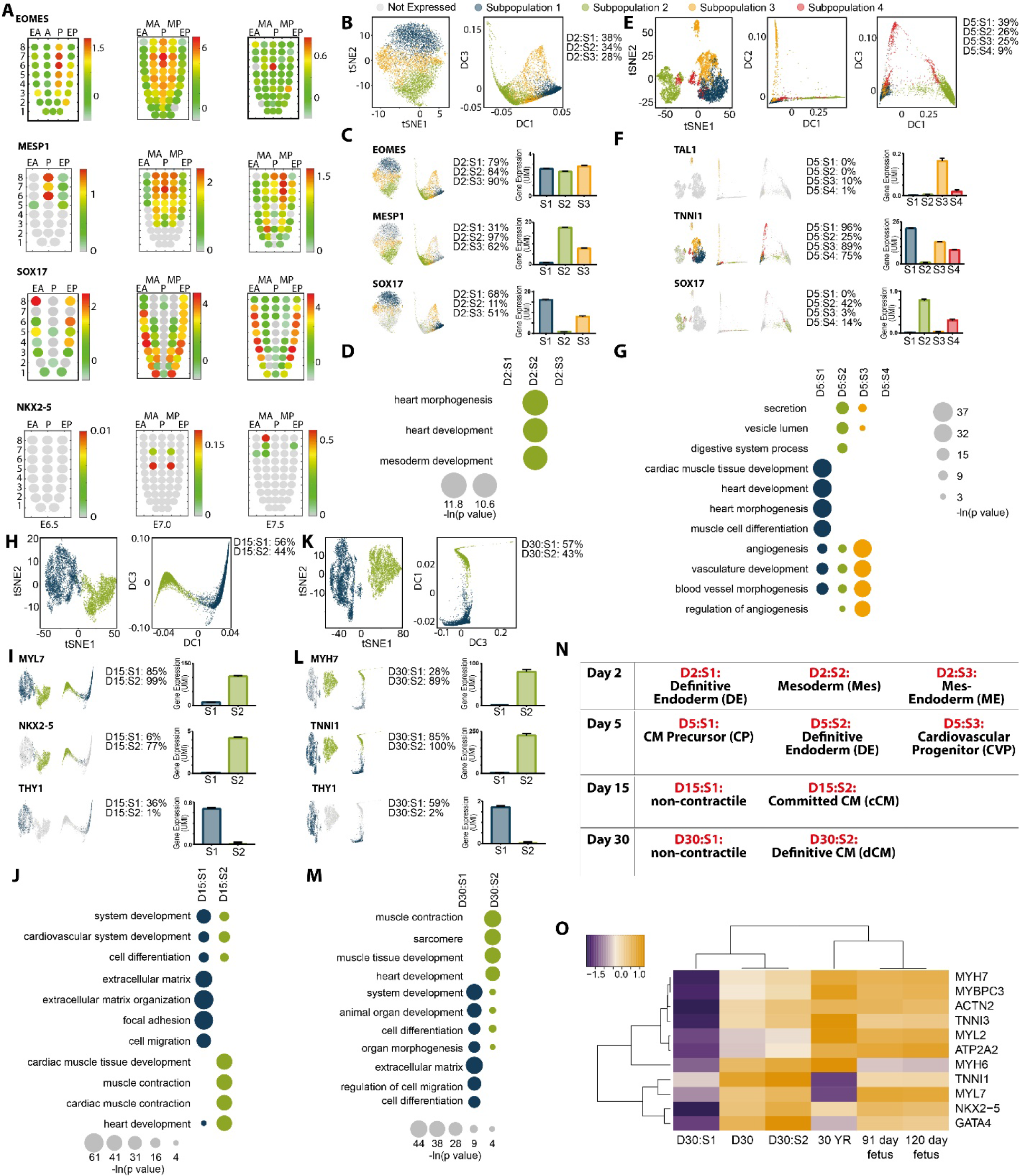
Subpopulation identification and characterization. **(A)** Corn plots showing spatial domains of EOMES, MESP1, SOX17 and NKX2-5 expression in the mesoderm and endoderm of E6.5, E7.0, and E7.5 mouse embryos during gastrulation (unpublished RNA-seq data for E6.5 (n = 6) and E7.5 (n = 6) embryos and published data for E7.0 embryo (Peng et al., 2016)). Positions of the cell populations (“kernels” in the 2D plot of RNA-Seq data) in the germ layers: the proximal-distal location in descending numerical order (1 = most distal site) and in the transverse plane of the mesoderm and endoderm - Anterior half (EA) and Posterior half (EP) of the endoderm, Anterior half (MA) and Posterior half (MP)of the mesoderm, and Posterior epiblast (P) containing the primitive streak. Color scales represent levels of expression as log10 of fragments per kilobase million (FPKM + 1) (see **Figure S3** for schematic of iTranscriptome). **(B-D)** Analysis of day 2 subpopulations represented by **(B**) Reference *t*-SNE (left) and diffusion (right) plots and the percent of cells in each subpopulations (D2:S1-S3), **(C)** analysis of primitive streak genes EOMES (pan-mesendoderm transcription factor), MESP1 (cardiogenic mesoderm transcription factor), and SOX17 (definitive endoderm transcription factor). Below each gene name are shown the following data from left to right: *t*-SNE plot and diffusion plot of cells expressing each gene, percent of cells expressing gene, expression level of gene in each subpopulation. **(D)** Gene ontology analysis of differentially expressed genes showing enrichment for networks governing cardiac development enriched in subpopulation 2. **(E-G)** Analysis of day 5 progenitor subpopulations represented by (E) reference *t*-SNE (left) and diffusion (right) plots and the percent of cells in each subpopulations (D5:S1-S4), **(F)** analysis of progenitor genes TAL1 (endothelial fate transcription factor), TNNI1 (early development sarcomere isoform of TNNI), and SOX17 (definitive endoderm transcription factor). Below each gene name are shown the following data from left to right: *t*-SNE plot and diffusion plot showing cells expressing each gene, percent of cells expressing gene, expression level of gene in each subpopulation. **(G)** Gene ontology analysis of differentially expressed genes showing enrichment for networks governing cardiac development (D5:S1), definitive endoderm (D5:S2), and endothelium (D5:S3). (H-J) Analysis of day 15 subpopulations represented by **(H)** reference *t*-SNE (left) and diffusion (right) plots and the percent of cells in each subpopulations (D15:S1-S2), **(I)** analysis of cardiac genes MYL7 (early development sarcomere isoform of MYL), NKX2-5 (cardiac transcription factor), and THY1 (fibroblast marker). Below each gene name are shown the following data from left to right: *t*-SNE plot and diffusion plot showing cells expressing each gene, percent of cells expressing gene, expression level of gene in each subpopulation. **(J)** Gene ontology analysis of differentially expressed genes showing enrichment for networks governing extracellular matrix and cell motility (D15:S1) and cardiac development (D15:S2). **(K-M)** Analysis of day 30 subpopulations represented by **(K)** reference t-SNE (left) and diffusion (right) plots and the percent of cells in each subpopulations (D30:S1-S2), **(L)** analysis of cardiac genes TNNI1 (early development sarcomere isoform of TNNI), MYH7 (mature sarcomere isoform of MYH), and THY1 (fibroblast marker). Below each gene name are shown the following data from left to right: t-SNE plot and diffusion plot showing cells expressing each gene, percent of cells expressing gene, expression level of gene in each subpopulation. **(M)** Gene ontology analysis of differentially expressed genes showing enrichment for networks governing system development and morphogenesis (D30:S1) and cardiac development (D30:S2). (N) Overall phenotypic determinations of subpopulation identity based on *in vivo* anchoring genes outlined through stage-specific transitions in differentiation. CM: cardiomyocyte. **(O)** Expression of cardiac genes in day 30 hPSC-derived cardiomyocytes (all cells vs S1 vs S2) relative to expression levels in human foetal and adult heart samples (ENCODE). Gene expression is measured as counts per million mapped reads and each gene is internally normalized to maximum expression. Subpopulation coloration across time points does not reflect developmental fate relationship. UMI: unique molecular identifier.

Based on these observations, we dissected the transcriptional phenotype of subpopulations identified during human cardiac directed differentiation. From pluripotency (**Figure S3A-B**), cells navigate through germ layer specification (day 2), comprising three transcriptionally distinct subpopulations that express the pan-mesendoderm gene, EOMES (**Figure 2B-C**, **Figure S2A**). Specific day 2 subpopulations express genes involved in mesoderm (D2:S2), mesendoderm (D2:S3), and definitive endoderm (D2:S1) (**Figure 2B-C** **and** **Figure S2D** **and** **Figure S3C-D**). Gene ontology (GO) analysis of differentially expressed genes between subpopulations indicated that only D2:S2 showed significant enrichment for cardiogenic gene networks (**Figure 2D**, **Table S1**). Surprisingly, these data show that only 34% of day 2 cells comprise cardiogenic mesoderm marked by MESP1 with the majority of cells characterized by mesendoderm and definitive endoderm expression patterns. At the progenitor stage (day 5), we identified cardiac precursors (D5:S1 and D5:S3) (**Figure 2E-G** **and** **Figure S3E**), a persistent population of definitive endoderm (D5:S2) (**Figure 2E-F** **and** **Figure S3E**), and endothelial cells (D5:S3) (**Figure 2E-G**). Day 15 and day 30 cells comprised two subpopulations (**Figure 2H-M** **and** **Figure S3F-G**). NKX2-5, MYH6, TTN and other cardiac structural and regulatory genes were identified in S2 (**Figure 2H-M** **and** **Figure S3F-G**). In contrast, S1 was primarily characterized by GO enrichment for genes associated with extracellular matrix deposition, motility, and cell adhesion (**Figure 2J** **and M**) which was supported by identification of a significant number of fibroblast-like cells marked by THY 1 (CD90) in S1 (**Figure 2I and L**). The co-existence of a non-contractile cell population, which is characterized as non-myocytes, is common in directed cardiac differentiation (Dubois et al., 2011). Taken together, these data show iPSC differentiation into committed (day 15) and definitive (day 30) cardiomyocytes (S2) and non-contractile cells (S1) (**Figure 2N**). To assess the level of maturity derived from this protocol relative to *in vivo* human development, we compared day 30 clusters against ENCODE RNA-seq data from foetal and adult hearts (**Figure 2O**). Using genes that reflect either early foetal (TNNI1, MYH6) vs late stages of heart development (MYH7, TNNI3, MYL2), the most differentiated *in vitro* derived cardiac population (D30:S2) remains more developmentally immature than even first trimester human hearts.

### Lineage predictions based on regulatory gene networks governing differentiation

While these bulk population analyses provided clarity into the diversity of cell types represented in cardiac differentiation, we sought to understand the lineage trajectories and gene regulatory networks governing diversification of cell fates. We implemented a probabilistic method for constructing regulatory networks from single-cell time series expression data (*scdiff*: Cell Differentiation Analysis Using Time-series Single-cell RNA-seq Data) (Ding et al., 2018). The algorithm utilizes TF-gene databases to model gene regulation relationships based on the directional changes in expression of TFs and target genes at parental and descendant states. These prerequisites impact both cell assignment and model learning since each state is represented by a probabilistic model that takes into account not just the expression but also the regulatory information. As such, *scdiff* does not exclusively rely on expression similarity to connect states allowing it to overcome problems related to sampling since it can still identify descendent states even if they are less similar in terms of their actual expression profiles.

We used *scdiff* to predict lineages underlying the diversity of fates during small molecule-mediated cardiac directed differentiation (**Table S2** and **Figure 3A**). Overall, the algorithm identified three distinct lineages from pluripotency comprising 10 nodes. Since this algorithm reassigns cells based on regulatory networks, we analyzed the distribution of cell subpopulations based on our CORE cluster classifications as outlined in **Figure 2** to establish population identities linking predicted lineages (**Figure 3A-B** **and** **Figure S4A**). The first lineage (N1:N2) diverts from pluripotency into a SOX17/FOXA2/EPCAM^+^ definitive endoderm population that terminates at day 2 and is comprised almost exclusively of D2:S1 and D2:S3 (**Figure 3A-B** **and** **Figure S4A**). The second lineage, N1:N3:N5, transitions from pluripotency (N1) into node 3 which is primarily comprised of definitive endoderm (D2:S1) and mesendoderm (D2:S3) but includes a larger fraction of MESP1/T^+^ mesoderm (D2:S2).This node is predicted to be the origin of another terminal lineage endpoint, node 5 at day 5, comprising FOXA2/EPCAM^+^ definitive endoderm cells (D5:S2 and D5:S4) (**Figure 3A-B** **and** **Figure S4A**). The third lineage comprises the longest trajectory through differentiation involving stepwise transitions in cardiac fate (N1:N4:N6-N9 and N6-N10). Pluripotent cells (N1) give rise initially to node 4 mesoderm (D2:S2) and mesendoderm (D2:S3) cells with subsequent progression into cardiac precursor cells (N6: primarily D5:S1 and D5:S3). From day 5 the algorithm predicts a bifurcation of fate giving rise to THY1^+^/NKX2-5^-^ non-contractile cardiac derivatives (N8-10: D15:S1 and D30:S1) or NKX2-5^+^/MYH6^+^ committed CM (N7: D15:S2) that progress onto MYH7^+^/MYL2^+^ definitive CM (N9: D30:S2) (**Figure 3A-B** **and** **Figure S4A**).

**Figure 3.**
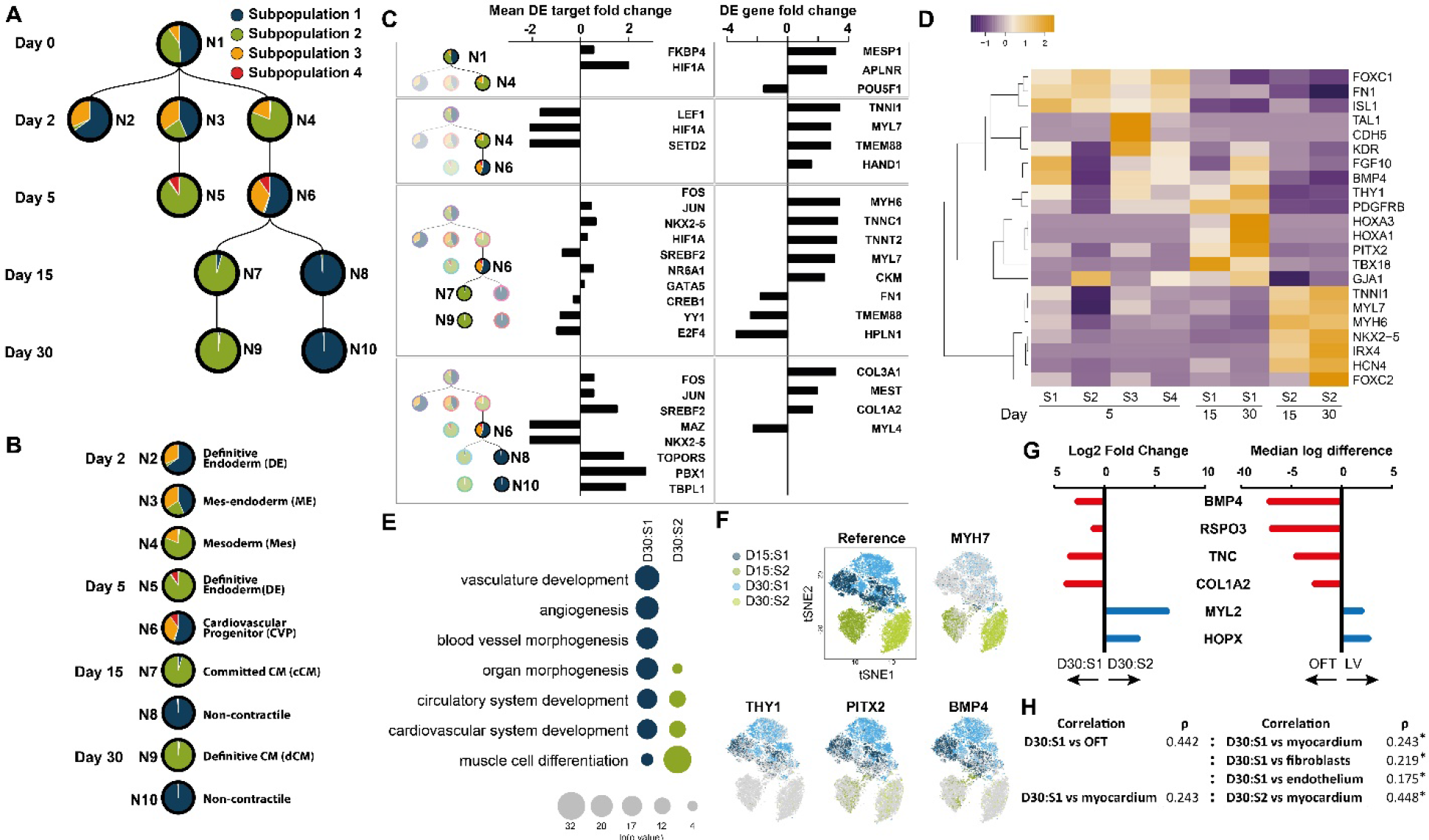
Transcription factor regulatory networks predict developmental fate choices during cardiac differentiation. **(A)** Stepwise transitions into cardiac lineages from pluripotency predicted on the basis of gene regulatory networks (GRN) detected between pairwise changes in cell state during differentiation. Circles indicate distinct nodes governed by a common GRN. Since cells can be reassigned based on the expression of their genes, the re-distribution of subpopulations established by clustering analysis and phenotyping as outlined in **Figure 2** are represented as pie charts within each circle indicating the percent of cells from each subpopulation contributing to that node. Each node is numbered N1-N10 for reference. **(B)** Phenotypic identity of nodes reflecting stage-specific transitions in cell state through cardiac directed differentiation. **(C)** Analysis of transcription factors (TFs) and genes controlling stage-specific regulatory networks underlying cell fate transitions. Mean DE target fold change calculates the fold change for the differentially expressed targets of the TF. DE gene fold change shows up or down-regulated fold change of TF target genes. **(D)** Heat map comparing expression across all cells from day 5, 15, and 30 subpopulations for genes involved in progenitor specification, vascular endothelial development, outflow tract development, and primary heart field cardiomyocyte development. Gene ontology analysis comparing day 30 S1 vs S2 showing gene networks involved in vascular development enriched in S1 vs cardiac muscle development enriched in S2. **(F)** *t*-SNE and diffusion plots for all cells from days 15 and 30 showing expression distribution of the cardiac gene MYH7 (high in S2 at day 15 and 30) relative to outflow tract development genes THY1, PITX2, and BMP4 (high in S1 at day 15 and 30). **(G)** The top most differentially expressed genes identified by *in vivo* single-cell analysis comparing outflow tract (OFT) vs ventricular cardiomyocyte (Li et al., 2016) were compared against their expression level in D30:S1 vs D30:S2 *in vitro* derived cardiac derivatives. **(H)** Differentially expressed genes between subpopulations D30:S1 and D30:S2 were used to assess transcriptional similarity to *in vivo* cell types using Spearman’s correlation analysis. Values are presented median Spearman’s value ρ. Significant differences between pairs of correlation coefficients were calculated using a Fisher Z-transformation. Subpopulation coloration across time points does not reflect developmental fate relationship. *P*-values for all tests were below the double precision limit of 2.2e-308

We leveraged the regulatory network predictions to identify key transcription factors and target genes underlying progressive fate changes across all 10 nodes (**Figure 3C** **and Table S2**). These data reinforce established mechanisms of cardiac lineage specification. In particular, we found evidence for down-regulation of Wnt/β-catenin signaling (LEF1) between N4-N6 which is required to transition from mesoderm into the cardiac progenitor cell (Paige et al., 2010; Palpant et al., 2013; Ueno et al., 2007).

From the progenitor node N6 into contractile cardiomyocytes N7:N9, the data show proper down-regulation of progenitor transcription factors such as YY1 and up-regulation of TFs known to control cardiomyocyte differentiation such as NKX2-5. Downstream target genes involved in governing the transition from progenitor cells to a differentiated cardiac state were expressed concomitantly (**Figure 3C**). Further subtype specification through computational analysis of D30:S2 did not reveal further atrial or ventricular cardiomyocyte subtypes by transcriptional profiling. Additionally, empirical studies are required to evaluate the composition, stoichiometry, and trajectory of these computationally determined lineages.

To gain new insights into lineage trajectories derived during differentiation, we sought to understand the gene network underlying specification of non-contractile cardiac derivatives N8:N10, a population currently not well defined although widely used for tissue engineering applications (Thavandiran et al., 2013). The predicted network underlying this transition showed significant down-regulation of cardiac TFs NKX2-5 and MAZ while other TFs involved in lipid metabolism/sterol regulation (SREBF2) and protein sumoylation (TOPORS) were up-regulated (**Figure 3C**). Of particular note, we observed up-regulation of Pre-B cell leukemia transcription homeobox (PBX1: *P* = 1.1e^-16^, mean DE target fold change = 2.72), a transcriptional regulator that activates a network of genes associated with cardiac outflow tract (OFT) morphogenesis (Arrington et al., 2012).

To further assess an OFT phenotype, we compared expression of a panel of cardiomyocyte, early developmental vascular endothelial, and OFT development genes across all subpopulations comprising transitions from day 5 to 30 (**Figure 3D**). While early developmental vascular EC differentiation genes (TAL1, CDH5) were expressed in D5:S3, these genes were not expressed in D15 or D30. Furthermore, while D15 and D30 S2 cells expressed cardiac sarcomere genes and transcription factors associated with first heart field specification (IRX4 and HCN4), S1 cells exclusively expressed an extensive network of genes associated with OFT development including PITX2, TBX18, HOXA1-3, FGF10, GJA1, and KDR (**Figure 3D**). We also performed gene ontology analysis of differentially expressed genes between D30:S1 (N10) and D30:S2 (N9) cells. These data show a significant enrichment for gene networks related to vascular development (*P* = 1.1e^−11^) and blood vessel morphogenesis (*P* = 4.7e^−9^) exclusively within node 10 D30:S1 cells. This finding is supported by single-cell visualization showing enrichment of OFT gene expression in S1 vs S2 by t-SNE analysis of THY1 (59% D30:S1 vs 2% D30:S2), BMP4 (70% D30:S1 vs 6% D30:S2), and PITX2 (73% D30:S1 vs 17% D30:S2) (**Figure 3E-F**).

Lastly, to anchor this observation to *in vivo* cell types, we used single-cell RNA-seq data of *in vivo* heart development (Li et al., 2016) to identify the top most differentially expressed genes between outflow tract and left ventricle (LV). These data show expression of BMP4, RSPO3, TNC, and COL1A2 in D30:S1 and *in vivo* OFT derivatives and MYL2 and HOPX upregulated in cardiomyocytes (**Figure 3G**). Lastly, to assess cell-type specific transcriptional signatures, we identified differentially expressed genes between D30 S1 vs S2 and performed a Spearman rank correlation analysis against expression profiles of *in vivo* FACS sorted (Quaife-Ryan et al., 2017) or single cell-derived cardiac subtypes (Li et al., 2016). These data show that D30:S1 has a significantly stronger correlation to OFT cells (Spearman’s ρ = 0.442) than fibroblasts (Spearman’s ρ = 0.219), endothelium (Spearman’s ρ = 0.175), or myocardium (Spearman’s ρ = 0.243) (*P* < 2.2×10^−16^ for all pairwise comparisons) (**Figure 3H**). As expected, there was a significantly stronger correlation of D30:S2 to myocardium (Spearman’s ρ = 0.448) compared to D30:S1 (Spearman’s ρ = 0.243, *P* < 2.2×10^−16^) (**Figure 3H**). Collectively, these data indicate that directed differentiation generates definitive cell populations comprising contractile cardiomyocytes and a non-contractile cell type whose transcriptional signature correlates with cardiac outflow tract cells. Due to the complex cellular origins of outflow tract and the diversity of non-contractile cell types of the heart *in vivo*, further studies are required to determine the specific identity and biology of these cells and their application in disease modelling and tissue engineering.

### HOPX is dysregulated during *in vitro* directed differentiation from hPSCs

Having established subpopulation identities and benchmarked *in vitro* derived subpopulations to *in vivo* development, we next aimed to identify dysregulated gene networks with the objective of determining novel mechanisms for modelling *in vitro* differentiation to more accurately reflect *in vivo* heart development. To this end, we analyzed a panel of 52 transcription factors and epigenetic regulators known to govern diversification of mesoderm and endoderm lineages represented in this data set (**Table S3**). Expression of these regulatory genes was measured across eleven subpopulations identified between days 2-30 of differentiation. HOPX, a non-DNA binding homeodomain protein identified in this analysis, has previously been shown to be one of the earliest, specific markers of cardiomyocyte development (Jain et al., 2015), and governs cardiac fate by regulating cardiac gene networks through interactions with transcription factors, epigenetic regulators, and signaling molecules Chen et al., 2002; Jain et al., 2015). We have also recently shown that HOPX functionally regulates blood formation from hemogenic endothelium (Palpant et al., 2017b). Consistent with mouse heart development, analysis of human foetal development at each trimester indicate a robust activation of HOPX during heart development *in vivo* (**Figure S5A**).

Previous studies have shown HOPX is expressed during cardiomyocyte specification at the progenitor stage of mouse development *in vivo* (Jain et al., 2015) whereas we detected HOPX only in endothelium (D5:C3) and not in cardiac precursor cells (D5:C1) at an equivalent time point (day 5) of *in vitro* differentiation (**Figure 4A-B**). Second, in contrast to previous studies *in vivo* where HOPX lineage traces almost all cardiomyocytes of the heart (Jain et al., 2015), HOPX is detected in only 16% of D30:S2 cardiomyocytes (**Figure 4B-D**). To rule out stochastic expression in cardiomyocytes due to low sequencing read depth resulting in dropout, we analyzed expression of a panel of genes known to regulate cardiac lineage specification and differentiation (**Figure 4C-D**). While HOPX is rarely detected, its expression level is equivalent to that of other cardiac TFs that are detected in a high percentage of D30:S2 cardiomyocytes (HAND1: 67%, HAND2: 64%, GATA4: 67%, NKX2-5: 86% vs HOPX: 16%) (**Figure 4B-D**).

**Figure 4.**
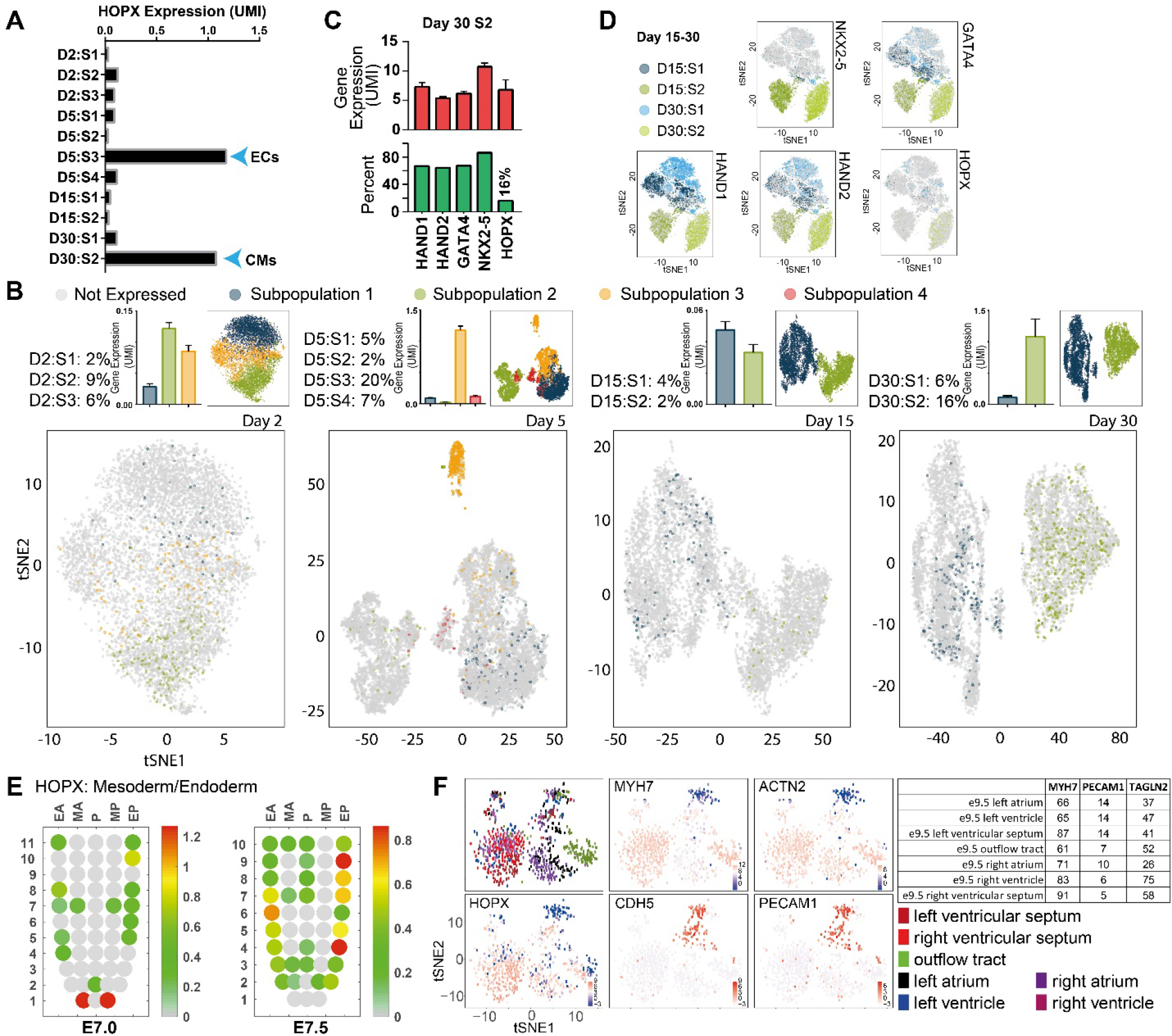
HOPX is rarely expressed during *in vitro* cardiac directed differentiation. **(A)** Analysis of HOPX expression in eleven subpopulations from day 2 to day 5 of differentiation showing expression as early as day 2 mesoderm and highest expression in day 5 endothelial cells (ECs) and day 30 cardiomyocytes (CMs). **(B)** Single-cell expression analysis of HOPX at day 2, 5, 15, and 30. Data presented include *t*-SNE plots indicating distribution and localization of HOPX expressing cells in different subpopulations (bottom), the percentage of HOPX^+^ cells in each subpopulation (top left), bar graphs showing expression of HOPX in each subpopulation (top middle), and the reference *t*-SNE plot demarcating subpopulations (top right). **(C)** Analysis of known genetic regulators of heart development only in subpopulation 2 at day 30 of differentiation. **(D)** *t*-SNE plots of merged data sets from two continuous days for all cells between day 15-30 for each gene showing robust distribution of key cardiac regulatory genes with the exception of HOPX. **(E)** Corn plots showing the spatial domains of HOPX expression in the mesoderm and endoderm of E7.0 and E7.5 mouse embryos during gastrulation (unpublished RNA-seq data for E7.5 (n = 6) embryos and published data for E7.0 embryo, (Peng et al., 2016)). Positions of the cell populations (“kernels” in the 2D plot of RNA-Seq data) and in the germ layers: the proximal-distal location in descending numerical order (1 = most distal site) in the transverse plane of the mesoderm and endoderm - Anterior half (EA) and Posterior half (EP) of the endoderm, Anterior half (MA) and Posterior half (MP) of the mesoderm, and Posterior epiblast (P) containing the primitive streak. Color scales represent levels of expression as log10 of fragments per kilobase million (FPKM+1). **(F)** Single-cell expression analysis of E9.5 mouse heart (Li et al., 2016) showing HOPX expression relative to markers of cardiomyocytes (MYH7, ACTN2) and endothelial cells (CDH5, PECAM1) (scale bars are Log2(RPM)). Table (right) shows percent of cardiac (MYH7), endothelial (PECAM1), and smooth muscle (TAGLN2) cells co-expressing HOPX in various regions of the developing mouse heart. UMI: unique molecular identifier.

We analyzed spatio-temporal gene expression throughout mesoderm and endoderm development in the gastrulating mouse *in vivo*. Assessment of gene expression during mouse gastrulation *in vivo* shows HOPX expression as early as E6.5 in the proximal portion of the nascent primitive streak (P) (**Figure 4E** and Figure S5B-C) similar to the expression pattern of MESP1 (**Figure S5D-E**). From E7.0 to E7.5, HOPX is increasingly expressed throughout the developing endoderm. By E7.5, HOPX displays residual expression in the remaining distal primitive streak, endoderm (EA to EP), and the anterior mesoderm (MA) in coordination with other cardiogenic genes including NKX2-5 and MESP1 (**Figure 4E** and **Figure 2A**). From E8.5 onward, cardiac development and morphogenesis occurs as chambers, valves, and outflow tract form. We analyzed HOPX expression across diverse cell types contributing to heart development *in vivo* using single-cell transcriptomic analysis of the E9.5 mouse heart (Li et al., 2016). These data indicate that HOPX expression is distributed throughout all chambers and cell types of the heart. While HOPX expression largely coincides with expression of cardiac genes MYH7 and ACTN2, HOPX is also expressed in endothelial cells (CDH5^+^ and/or PECAM1^+^), smooth muscle cells (MYH11^+^ and/or TAGLIN2^+^), and epicardial cells (WT1^+^) (**Figure 4F** **and Table S4**).

### Lineage trajectory of HOPX-expressing cells *in vitro*

We analyzed the lineage trajectory of HOPX^+^ cells at single-cell resolution during cardiac directed differentiation to determine the core gene networks and transcription factors governing successive fate choices of HOPX expressing cells during cardiac differentiation (**Figure S4B-D**, **and Table S5**). At day 2 rare HOPX expressing cells are identified in mes-endoderm (D2:S2 9% and D2:S3 6%) and rarely in definitive endoderm (D2:S1 2%) with the HOPX lineage at this early stage of specification comprising 2 lineages (N2 and N3) enriched for expression of cardiogenic mesoderm genes such as MESP1 (**Figure S4C-D**). From day 2 into day 5, HOPX^+^ cells remain sparse in progenitor cell populations where day 2 N3 splits into two day 5 lineages N4 and N5 (**Figure S4B**). Based on lineage prediction, an equal proportion of HOPX^+^ cells give rise to TNNI1^+^ cardiac precursor cells (N4: 389 cells) or TAL1^+^ expressing endothelial cells (N5: 381 cells) with both fates governed by established TFs and downstream gene networks required for endothelial (NRP2, KDR) vs cardiac fate specification (TNNI1, TMEM88) (Figure S4C-D and Table S5). Progressing to day 15 of differentiation, HOPX remains rare (2-4% of cells) and splits into two separate lineages derived from day 5 cardiac precursor cells (N4). Governed in part by increased NKX2-5 and downregulation of the cardiac progenitor TF YY1, HOPX cardiac precursor cells differentiate into MYL2/IRX4^+^ cardiomyocytes (N6-N7) while a separate branch governed by TFs such as PBX1 differentiate into non-contractile derivatives (N8-N9) (**Figure S4B-D**). Overall, analysis of HOPX gene expression across populations shows that HOPX is most highly expressed in endothelial cells at day 5 and in committed cardiomyocytes at day 30 of directed differentiation (**Figure 4A**).

### Chromatin and expression analysis of HOPX in cardiac lineage specification

To determine the epigenetic basis for HOPX dysregulation during *in vitro* differentiation, we analyzed chromatin and transcriptional regulation at the HOPX locus (Palpant et al., 2017a) (**Figure S5F**). Chromatin immunoprecipitation data for repressive chromatin (H3K27me3), actively transcribed chromatin (H3K4me3), and gene expression by RNA-seq (Palpant et al., 2017b) show that in the context of cardiac directed differentiation HOPX is epigenetically repressed on the basis of abundant H3K27me3 compared to H3K4me3 in day 5 cardiac precursor cells (**Figure S5F**). This is consistent with RNA-seq, qRT-PCR, and analysis of HOPX activity in knockin HOPX reporter cells showing that HOPX is expressed late during cardiac differentiation, well after sarcomere formation and weeks after the onset of spontaneously beating cells during cardiac directed differentiation (**Figures 4B**, **S5F, and S6A-D**). The highest level of HOPX expression was observed in cardiomyocyte cultures maintained for 1 year (**Figure S6F-G**). Collectively, these data show a direct link between chromatin regulation of the HOPX locus and expression of HOPX in cardiac lineage specification *in vitro*.

### HOPX drives cardiomyocyte hypertrophy

To determine the functional role of HOPX in *in* vitro-derived cardiomyocytes, we established conditional HOPX over-expression hPSCs in which a nuclear localized HOPX is targeted to the AAVS1 locus (**Figure 5A** **and** **Figure S7**). Using western blot, qRT-PCR, and immunostaining, we show that HOPX is over-expressed in a doxycycline inducible manner and is nuclear localized (****Figure 5A-C****). Morphometric analysis of dox treated cardiomyocytes showed a significant increase in cell area under conditions of HOPX overexpression (**Figure 5D**). We performed bulk RNA sequencing analysis on control vs HOPX OE cardiomyocytes to determine global transcriptomic changes (**Figure 5E-H** **and Table S6**). Analysis of differentially expressed genes showed a significant enrichment of gene ontologies associated with signaling pathways (ERK1-2, IGF) and gene networks involved in cell growth and maturation in HOPX OE cardiomyocytes with IGF-1 representing the most highly up-regulated among a panel of known regulators of hypertrophy (**Figure 5G-H** **and Table S7**).

**Figure 5.**
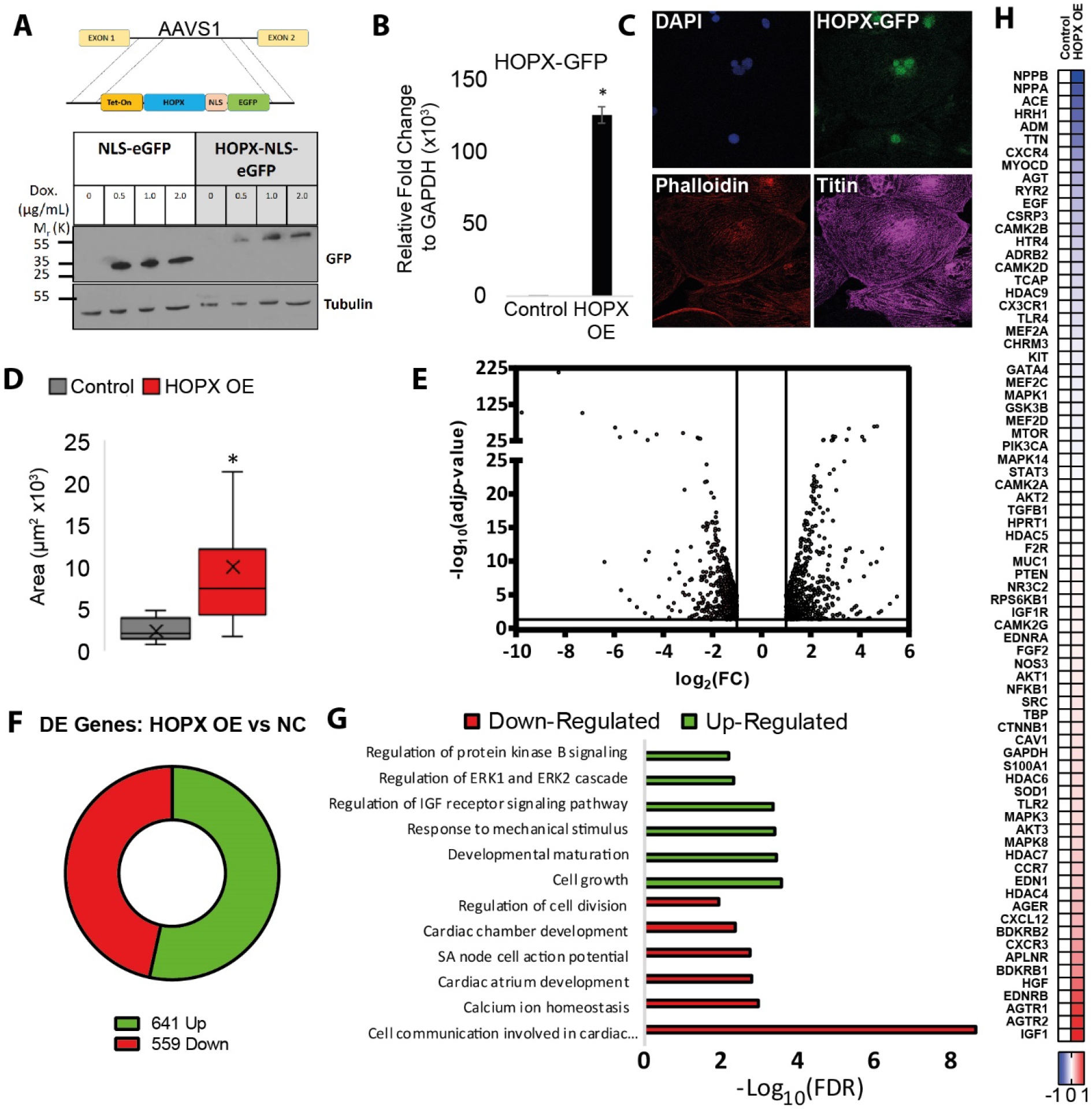
HOPX is a key regulator of cardiomyocyte hypertrophy. **(A)** Gene targeting strategy for conditional HOPX over-expression. Schematic shows design of conditionally expressed HOPX-NLS-EGFP construct. Below, western showing doxycycline induction of control (NLS-eGFP) and HOPX OE iPSC lines. **(B)** Quantitative PCR analysis of HOPX transcript abundance in control vs HOPX OE iPSCs. **(C)** Immunohistochemistry showing nuclear localization of HOPX-GFP in cardiomyocytes. **(D)** Cell size analysis showed that HOPX OE treated hiPSC-CMs led to a significant increase in area. **(E-G)** Volcano plot (E), quantification of DE genes **(F)**, and gene ontology analysis (G) of differentially expressed genes identified by RNA-seq of control vs HOPX OE cardiomyocytes. **(H)** Genes known to govern hypertrophy showing IGF1 as the most significantly upregulated hypertrophic gene in HOPX OE vs control cells. For heat maps, data are presented as Log10 transformed relative gene expression normalized to HPRT. NLS: nuclear localization signal, EGFP: enhanced green fluorescent protein. Scale bars = 100 μm. * P <0.05 by t test.

### The HOPX locus is activated by hypertrophic stimulation

These over-expression data indicate that HOPX is sufficient to drive cardiomyocyte hypertrophy *in vitro*. Hypertrophic stimulation is not modelled well in high density monolayer cardiac directed due to the absence of mechanical stretch or exogenous hypertrophic signals present during *in vivo* heart development. On the basis that this may, at least in part, explain the dysregulation of HOPX *in vitro*, we tested the reciprocal hypothesis: whether exogenous hypertrophic stimuli is sufficient to drive HOPX expression *in vitro*. To this end, we implemented an established approach for stimulating hypertrophy (Uesugi et al., 2014) in which high-density monolayer-derived cardiomyocytes are replated at low density at day 10 and analyzed at day 15 (**Figure 6A**). In keeping with a hypertrophic response, replating cardiomyocytes results in significantly increased cell area and anisotropy (**Figure 6B-C**) and up-regulation of genes known to govern hypertrophic growth in cardiomyocytes including NPPB, MYOCD, EDN1, IGF1 and others (**Figure 6D**). Importantly, we found by quantitative PCR analysis that replating cardiomyocytes resulted in a greater than 10 fold increase in HOPX (**Figure 6E**) in coordination with significant increases in expression of myofibrillar isoforms (MYH7, MYL2, TNNI3) and transcription factors (SRF) involved in cardiomyocyte maturation (**Figure 6F**). To assess HOPX expression at single cell resolution, we utilized HOPX-reporter hPSCs in which tdTomato is knocked into the translational start site of HOPX (Palpant et al., 2017b) (**Figure S6C**). These data show that HOPX is robustly activated uniformly in replated cardiomyocytes (**Figure 6G**). We further show that treatment with Endothelin-1 (ET-1), a potent stimulus of cardiomyocyte hypertrophy, significantly increases HOPX expression albeit to a much lower level than replated cardiomyocytes (**Figure 6H**). Taken together, these results indicate that transcriptional activation of the HOPX locus is downstream of hypertrophic signaling.

**Figure 6.**
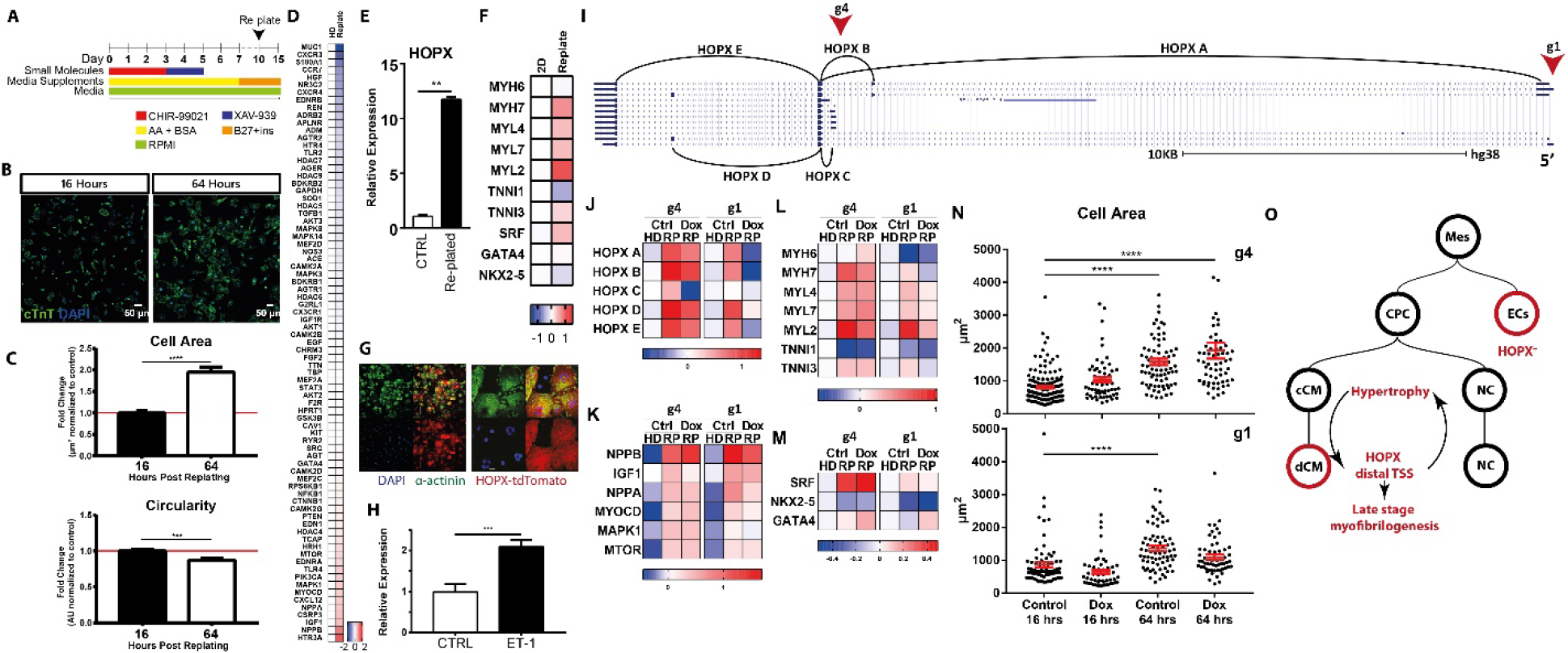
HOPX functionally governs cardiac hypertrophy through the distal transcriptional start site. **(A)** Schematic of *in vitro* directed differentiation of hPSCs with re-plating at day 10 and analysis at day 15. **(B-C)** Representative images **(B)** and quantification **(C)** of morphometric changes during replating including cell area and circularity. **(D)** Quantitative PCR analysis of a selected panel of hypertrophic genes differentially expressed in the context of replating cardiomyocytes. HD: High density monolayer. **(E-F)** Quantative PCR analysis showing significant increases in HOPX **(E)** among a range of other cardiac transcription factors and myofilament genes **(F)** involved in cardiomyocyte maturation in replated cardiomyocytes compared to controls. **(G)** Immunohistochemistry of HOPX-tdTomato reporter cells showing uniform expression of HOPX in α-actinin^+^ replated cardiomyocytes. **(H)** Treatment with the hypertrophic signaling molecule Endothelin-1 (ET1) significantly increases HOPX expression. **(I)** UCSC genome browser analysis of transcript variants mapped to the HOPX locus, the position of guide RNAs blocking the proximal (g1) or distal (g4) TSS, and position of qPCR primers amplifying different exons of the HOPX locus. **(J-M)** Analysis of gene expression in control high density monolayer cells vs replated cells and HOPX KD replated cells by quantitative PCR for various exons of the HOPX locus as outlined in panel H **(J)**, genes governing cardiomyocyte hypertrophy **(K)**, cardiac myofilament genes **(L)**, and cardiac transcription factors **(M). (N)** Morphometric analysis of cell area in control vs HOPX KD cells over 64 hrs of replating. (O) Schematic lineage tree showing fate choices governed by HOPX during cardiac directed differentiation and a proposed mechanism whereby hypertrophic signaling is identified as a stimulus required for expression of HOPX during *in vitro* differentiation and showing that HOPX engages with cardiomyocyte hypertrophic growth through its distal transcriptional start site. For heat maps, data are presented as Log10 transformed relative gene expression normalized to HPRT. * *P* < 0.05. Data are presented as mean ± SEM.

### Dissecting the transcriptional complexity of HOPX regulation underlying cardiomyocyte hypertrophy

While genetic loss of HOPX did not impact specification of cardiomyocytes (**Figure S6E-F**), we set out to study the functional requirement and underlying complexity of the HOPX locus in cardiomyocyte hypertrophy. To this end, we utilized CRISPRi loss of function hPSCs to conditionally block HOPX expression at each of its two transcriptional start sites which we term the proximal TSS (inhibited by guide 4, g4) and distal TSS (inhibited by guide 1, g1) relative to the HOPX translational start site (**Figure 6I**). Quantitative PCR primers were designed to amplify different exons of the HOPX locus to map the transcriptional landscape of HOPX relative to the distal or proximal TSS in the context of hypertrophic stimulation (**Figure 6I**, **Table S8**). Cells were differentiated into the cardiac lineage +/− dox and analyzed at day 15 of differentiation under standard high density monolayer conditions vs replating. Transcriptional analysis of HOPX revealed a striking difference in the regulation of HOPX expression in the context of hypertrophy. As expected, all HOPX transcripts were significantly increased during replating (**Figure 6J**, **Table S8**). However, inhibition of the proximal TSS (g4) repressed expression from that locus (HOPX C) with no effect on transcriptional activity from the distal TSS (HOPX A) (**Figure 6J**). In contrast, inhibition of the distal HOPX TSS (g1) resulted in a global reduction of HOPX expression (HOPX A-E) with no transcriptional compensation from the proximal TSS (HOPX C) (**Figure 6J**). This indicates that HOPX has functionally distinct transcriptional start sites with the distal TSS functioning as the primary target of regulatory factors driving expression of HOPX in the context of hypertrophic stimulation.

To further determine the functional requirement of the proximal or distal HOPX TSS in hypertrophy, we analyzed hypertrophy-related genes up-regulated in replating (**Figure 6K**, **Table S8**). Loss of HOPX function from the distal or proximal TSS did not impact expression of any hypertrophic genes tested including IGF1 (**Figure 6K**), the most highly upregulated hypertrophy gene in HOPX OE (**Figure 5I**). We next assessed a panel of cardiac myofilament genes and transcription factors to determine the downstream impact on genes associated with cardiomyocyte maturation. We found that myofibrillar genetic isoforms associated with late stages of cardiomyocyte maturation (MYL2, MYH6, MYH7, TNNI3) were significantly depleted when the distal HOPX TSS was inhibited (g1). However, isoforms associated with early cardiac development (MYL4, MYL7, TNNI1) were not impacted. In contrast, knockdown of the proximal TSS (g4) had small but significant effects on expression of MYL2 and MYH7 but a significant increase in expression of the early fetal MYH6 isoform relative to controls (**Figure 6L**, **Table S8**). Expression of selected key cardiac transcription factors were not impacted by HOPX loss of function from either TSS with the exception of a significant increase in GATA4 with knockdown of the distal TSS (g1) (**Figure 6M**, **Table S8**). These data indicate that while genetic networks underlying hypertrophy and early cardiomyocyte myofibrillar development are not dependent on HOPX, we found it plays a key role driving maturation at least in part through regulating expression of late-stage genetic isoforms of myofibrillogenesis with the distal TSS playing a more dominant role compared to the proximal TSS.

We next assessed the impact of HOPX loss of function on morphometric parameters of cardiomyocyte hypertrophy. Cell area was measured at two time points post replating and showed a progressive and significant increase in cell size indicative of cardiomyocyte cell hypertrophy and maturation in control cells and conditions blocking the proximal HOPX TSS (g4) (**Figure 6N**). However, blocking transcription of the distal HOPX TSS attenuated the hypertrophic growth (**Figure 6N**).

Taken together, we have identified HOPX is a known key epigenetic regulator of cardiovascular development that is dysregulated during *in vitro* directed differentiation from hPSCs. Lineage tracing and transcriptional analysis of single cells transiting cardiac differentiation show HOPX predominantly expressed in day 5 endothelium and day 30 definitive cardiomyocytes (**Figure 6O**). We used genetic gain and loss of function models to determine mechanisms underlying transcriptional regulation of HOPX with the aim of directing gene networks underlying *in vitro* differentiation more closely mimicking *in vivo* development. We show that HOPX is situated downstream from hypertrophic signaling pathways, a stimulus not mimicked effectively in high density monolayer differentiation. Furthermore, HOPX activation directly drives hypertrophic growth and is essential for downstream expression of cardiac myofibrillar genetic isoforms involved in cardiomyocyte cell growth and maturation. Through genetic dissection underlying the transcriptional landscape of HOPX, we show that the distal HOPX TSS is the primary regulatory driver of HOPX activity underlying hypertrophic stimulation (**Figure 6O**).

## Discussion

This study provides single-cell resolution RNA-sequencing of human cardiac directed *in vitro* differentiation. Transcriptomic analysis of 43,168 cells traversing stepwise transitions in fate revealed cellular heterogeneity and the underlying gene networks involved in cardiac fate choices from pluripotency. The identification and characterization of *in vitro* derived cell types are supported by spatio-temporal gene expression of the gastrulating mouse embryo and single-cell analysis of *in vivo* heart development, providing a direct link to the complex restriction of fates underlying cardiovascular lineage specification *in vivo*. We leverage the computational power of single-cell level analysis to identify the framework of transcription factors and gene regulatory networks underlying progressive diversification of fates from pluripotency into mesoderm, definitive endoderm, endothelium, cardiomyocytes, and outflow tract fates. Transcriptional analysis of subpopulations revealed context-specific functions of transcription factors and epigenetic regulators underlying cardiovascular fate choices. We utilize this resource to identify novel mechanisms driving *in vitro* differentiation more accurately reflecting *in vivo* heart development and reveal the complex transcriptional landscape of HOPX in regulating cardiomyocyte hypertrophy. Collectively, we use a widely implemented protocol using Wnt modulation for cardiac directed differentiation (Burridge et al., 2014; Lian et al., 2012) with CRISPRi hiPSCs (Mandegar et al., 2016) providing a platform to dissect cardiac differentiation at single-cell resolution.

While the progression of heart development and morphogenesis from time points spanning E8.5 to P21 have been analyzed at single-cell resolution (DeLaughter et al., 2016; Li et al., 2016), a comprehensive transcriptomic profiling of the lineages derived by human pluripotent stem cell directed differentiation from pluripotency has not been available. Cardiac directed differentiation protocols using small molecules to modulate Wnt signaling have emerged in recent years as a simple, cost-effective, and reliable method to generate high-purity cardiac derivatives from hPSCs. Stem cell-derived cardiomyocytes generated using this approach have been utilized in translational applications to model patient-specific diseases (Ang et al., 2016; Bayzigitov et al., 2016; Ebert et al., 2014; Smith et al., 2017; Wu et al., 2015), test cardiotoxicity (Maillet et al., 2016), screen novel therapeutic drugs (Casini et al., 2017; Sharma et al., 2014), generate engineered heart tissue constructs to model the 3D environment of the heart (Huebsch et al., 2016; Tzatzalos et al., 2016), and develop cell-based regenerative therapies to repair heart tissue post-infarct (Hartman et al., 2016). In light of these diverse applications of this protocol in cardiovascular discovery and translational research, the current study provides whole genome-wide analysis of 43,168 transcriptomes undergoing stage-specific changes in gene expression during cardiac differentiation as a resource with which to dissect cell subpopulations at the molecular level.

Analysis of subpopulations during early stages of differentiation indicate a surprising contribution of mesendoderm and definitive endoderm coordinately specified with cardiac fates through the progenitor stage of differentiation. In particular, a minority of cells (34%) comprise MESP1^+^ cardiogenic mesoderm at day 2 that ultimately give rise to all cardiac derivatives at day 30. The interaction between endoderm and mesoderm in governing lineage specification *in vivo* is well known, and these data suggest that a critical functional role of induction cues provided by directed differentiation protocols is to establish the necessary population stoichiometry of transiently sustained endoderm required to support mesoderm in the derivation of high purity cardiac fates *in vitro*.

We evaluated lineage trajectories from single-cell data by implementing a lineage prediction algorithm, *scdiff*, specifically designed for learning regulatory networks controlling differentiation from single-cell time series data. This method relies on iterative analysis of known transcription factor-gene interactions to establish a gene regulatory framework that statistically links disparate populations captured across intermittent time intervals through differentiation. Analysis of cardiac directed differentiation using *scdiff* revealed stage-specific genetic regulators underlying diversification of cardiac fate from pluripotency. In particular, these data revealed new insights into the bifurcation of cardiac precursor cells at day 5 of differentiation into NKX2-5^+^/MYL2^+^ ventricular cardiomyocytes and a population of non-contractile cells with transcriptional networks similar to NKX2-5^−^/PITX2^+^ cardiac outflow tract (OFT) cells. While previous studies have routinely described a non-contractile THY1^+^ (CD90^+^) fibroblast-like cell used commonly for tissue engineering applications (Dubois et al., 2011; Thavandiran et al., 2013), this population remains poorly studied with no strong evidence for an *in vivo* correlate. Using transcriptional fate mapping and gene network analysis, we provide single-cell level transcriptome-wide evidence directly linked to *in vivo* cell cardiac types that non-contractile THY1^+^ cells are similar to cardiac OFT derivatives. Of importance, congenital heart disease (CHD) is among the most common forms of congenital defects (van der Linde et al., 2011), and OFT anomalies account for roughly 30% of CHD incidences (Thom et al., 2006). Given the complexity of outflow tract differentiation and morphogenesis that involves cell types form diverse origins, future work will require analysis of this population as it pertains to the cellular origins of outflow tract, septum, and other non-contractile cell types of the heart. While the genetic basis of OFT and septal malformations has been well studied (Arrington et al., 2012), the capacity to study development and disease using hPSC models has not been possible. Our findings have provided new insights into a cell subpopulation derived from *in vitro* cardiac directed differentiation that present new opportunities to develop translational platforms utilizing this cell type for disease modelling or therapies.

It is well-established that *in vitro* cardiac differentiation does not generate cardiomyocytes with the transcriptional profile, cellular diversity, morphometry, or functional maturity of adult *in* vivo-derived cardiomyocytes (Yang et al., 2014). This is at least in part a consequence of dysregulation of the stage-specific gene networks not properly modelled *in vitro*. In this study, we analyzed a panel of 52 transcription factors and epigenetic regulators across eleven cardiac differentiation-derived cell subpopulations at single-cell resolution. While HOPX is a key developmental regulator of cardiac myoblasts early in heart development *in vivo* (Jain et al., 2015) and data from this and other studies (Chen et al., 2002; Trivedi et al., 2010) showing a key role in heart maturation, we observe it rarely expressed in *in vitro* derived cardiomyocytes. We tested the hypothesis that the dysregulation of HOPX was the consequence of deficiencies in directed differentiation accurately mimic the signaling and mechanical stimuli of the developing heart. To address this, we aimed to understand the basis for activating HOPX and its downstream gene networks *in vitro* by identifying the upstream cues required for its expression. Utilizing gain and loss of function genetic models, we provide a comprehensive profiling of the complex transcriptional landscape of HOPX as a central regulator of the cardiomyocyte response to hypertrophy. These data show that the distal TSS is the primary hypertrophy responsive element and regulation of HOPX through this TSS functionally governs gene networks and cellular morphometric growth associated with cardiomyocyte hypertrophy and maturation.

Taken together, our study provides transcriptional profiling of human *in vitro* cardiac differentiation from more than forty thousand single-cells revealing cell diversity and genetic networks governing lineage progression of hPSC cardiac directed differentiation by small molecule Wnt modulation. These data provide new insights into the complexity of cell populations represented in stage-specific transitions from pluripotency and, coupled with the use of CRISPRi loss of function, establish a unique reference point for dissecting gene networks involved in human cardiac development and disease. Promoting adult-like phenotypes from *in vitro* differentiated cell types is essential for the realization of the translational applications of hPSCs in disease modelling and therapy. This study provides evidence that HOPX expression is a key transcriptional regulator near absent during high density cardiac directed differentiation *in vitro*, requiring hypertrophic stimulation to accurately direct HOPX and its downstream networks underlying the transcriptional and functional maturity of hPSC-derived cardiomyocytes.

## Acknowledgements.

Sequencing was performed by the Institute for Molecular Bioscience Sequencing Facility at the University of Queensland. Assistance with **Figure 1A** schematic was provided by Suzy Hur. The WTC CRISPRi hiPSCs and pQM plasmid backbone were kindly provided by the Conklin lab (UCSF, Gladstone Institute). We thank Prof Richard Harvey (Victor Change Cardiac Research Institute) for assistance reviewing the draft manuscript. This work was supported by the Australian Research Council (SR1101002) (NJP), the ARC Discovery Early Career Award (DE160100755) (ESW), National Health and Medical Research Council grants 1107599 and 1083405 (JP). ZBJ and JD were supported in part by grant 1R01GM122096 from the National Institute of Health, USA. This work was also supported by a Strategic Priority Research Program of the Chinese Academy of Sciences (XDA01010201 to N.J., XDA01010303 to J.D.J.H), National Key Basic Research and Development Program of China (2014CB964804, 2015CB964500, 2015CB964803), and National Natural Science Foundation of China (91219303, 31430058, 31401261, 91329302, 31210103916, and 91519330). P.P.L.T. is a Senior Principal Research Fellow of the National Health and Medical Research Council of Australia (1110751).

## Author contributions

**CEF:** Generated cells for single-cell RNA-seq, performed cell-based experiments including HOPX LOF, and wrote the manuscript

**QN:** Primary lead on computational analysis of single-cell data and wrote the manuscript

**SWL:** Performed single-cell isolation, barcoding, and sequencing and performed computational analysis of single-cell RNA-seq data and edited the manuscript

**AH:** Performed computational analysis of single-cell RNA-seq data and edited the manuscript HSC: Generated cells for single-cell RNA-seq

**JM:** Generated the HOPX inducible over-expression cell line and carried out HOPX OE assays

**SSS:** Conceived and generated iTranscriptome data for mouse gastrulation *in vivo*

**JDJH:** Conceived and generated iTranscriptome data for mouse gastrulation *in vivo*

**OP:** Assisted with the preparation of iTranscriptome data for publication

**GP:** Conceived and generated iTranscriptome data for mouse gastrulation *in vivo*

**GJB:** Assisted with sequencing

**AS:** Performed computational analysis of single-cell data ANC: Assisted with sequencing TJB: Assisted with sequencing

**NJ:** Conceived and generated iTranscriptome data for mouse gastrulation *in vivo*

**CEM:** Supervised HOPX LOF experiments and edited the manuscript

**ESW:** Adapted *scdiff* for large scale single-cell data sets and generated *scdiff* data

**JD:** Conceived of and developed *scdiff*

**ZBJ:** Conceived of and developed *scdiff* and wrote the manuscript

**HRB:** Supervised HOPX over-expression assays

**YW:** Performed computational analysis of single-cell data and edited the manuscript

**JH:** Assisted with bioengineered heart tissue experiments and edited the manuscript PPLT: Supervised iTranscriptome analysis, consulted single-cell phenotypes and edited the manuscript

**JEP:** Conceived and supervised experiments involving computational genomics analysis of single-cell RNA-seq data, wrote manuscript, and generated funding for the work

**NJP:** Conceived and supervised experiments involving stem cell differentiation, performed HOPX experiments, wrote manuscript, and generated funding for the work.

**See Supplemental Information for extended methods.**

## Cardiac directed differentiation using small molecule Wnt modulation at single-cell resolution

### Supplemental Information

#### Methods

##### Cell Culture

All human pluripotent stem cell studies were carried out in accordance with consent from the University of Queensland’s Institutional Human Research Ethics approval (HREC#: 2015001434). Human RUES2 hESCs and WTC11 hiPSCs were maintained on Vitronectin XF (Stem Cell Technologies, 07180) coated plates in mTeSR media (Stem Cell Technologies, 05850). Unless otherwise specified, cardiomyocyte directed differentiation using a monolayer platform was performed with a modified protocol based on previous reports (Burridge et al., 2014; Lian et al., 2012). On day −1 of differentiation, hPSCs were dissociated using 0.5% EDTA, plated into vitronectin coated plates at a density of 1.8 × 10^5^ cells/cm^2^, and cultured overnight in mTeSR media. Differentiation was induced on day 0 by changing the culture media to RPMI (Life Technologies Australia, 11875119) containing 3μM CHIR-99021 (Stem Cell Technologies, 72054), 500μg/mL BSA (Sigma Aldrich, A9418), and 213μg/mL ascorbic acid (Sigma Aldrich, A8960). After 3 days of culture, the media was replaced with RPMI containing 500μg/mL BSA, 213μg/mL ascorbic acid and 1μM Xav-939 (Stem Cell Technologies, 72674). On day 5, the media was exchanged for RPMI containing 500μg/mL BSA, and 213μg/mL ascorbic acid without supplemental cytokines. From day 7 onwards, the cultures were fed every 2 days with RPMI plus 1x B27 supplement plus insulin (Life Technologies Australia, 17504001). For endothelin 1 assays, cells were treated with 300 nM ET-1 (Sigma-Aldrich, E7764) from day 9-15 of directed differentiation. For HOPX over-expression studies (**Figure 5**), the following protocol was utilized: A monolayer-based directed differentiation protocol was followed to generate hiPSC-CMs, as described previously (Palpant et al., 2017a). On day 15 hiPSC-CMs were enriched by lactate selection (Tohyama et al., 2013).

##### Genetically modified cell lines

HOPX reporter cells were generated as previously described (Palpant et al., 2017b). WTC CRISPRi cells were generously provided by Bruce Conklin (UCSF, Gladstone Institute) as previously described (Mandegar et al., 2016). For HOPX loss of function studies, WTC CRISPRi hiPSCs were used. HOPX-targeted guide RNAs (gRNA) (**Table S9**) were designed to target sequences near the human HOPX distal and proximal transcription start sites, were cloned into the pQM-u6g-CNKB doxycycline-inducible construct and transfected into WTC CRISPRi GCaMP hiPSCs using GeneJuice Transfection Reagent (Merck, 70967). Stable clones were selected using successive rounds of re-plating with blasticidin (10 μg/ml). Populations were tested for knockdown efficiency by qPCR following doxycycline addition continuously from day 0 of cardiac-directed differentiation. HOPX OE-line creation: 1×10^6^ WTC hiPSCs were transfected with 0.5μg AAVS1-TALEN, 0.5μg AAVS1-TALEN and 4μg of HOPX-NLS-eGFP or 4μg of NLS-eGFP to generate the HOPX OE line and the HOPX negative control (NC) line using Amaxa Lonza Human stem cell Kit Ȳ2 (**Figure S7**). The cells were then plated with 5mM Rocki onto matrigel in mTeSR. Two days following the nucleofection, the cells were selected for Puromycin 0.5μg/ml for 2 days. The sequence for the HOPX-NLS-eGFP construct can be found in **Figure S7**.

##### Quantitative RT-PCR

For quantitative RT-PCR, total RNA was isolated using the RNeasy Mini kit (Qiagen, 74106). First-strand cDNA synthesis was generated using the Superscript III First Strand Synthesis System (Life Technologies Australia, 18080051). Quantitative RT-PCR was performed using SYBR Green PCR Master Mix (Life Technologies Australia, 4312704) on a ViiA 7 Real-Time PCR System (Applied Biosystems). The copy number for each transcript is expressed relative to that of housekeeping gene HPRT. Primers used for quantitative PCR are listed in **Table S6**. Quantification of cardiac hypertrophy gene expression was performed using Cardiac Hypertrophy H384 qPCR panels (BioRad, 10025144) with SYBR Green PCR Master Mix. Samples were run in biological triplicate. The copy number for each transcript is expressed relative to that of housekeeping gene HPRT1. FC was calculated on a gene by gene basis as gene expression divided by control gene expression.

##### Immunofluorescence and Morphometric analysis

Cells were fixed with 4% paraformaldehyde, permeabilized in PBS containing 0.025% Triton-X, and blocked in PBS containing 1.5% normal goat serum. Cells were stained with alpha-actinin (Sigma, Clone EA-53; Cat.#A7811, 1:800) and dsRed (Clontech, 632496) followed by secondary staining with AlexaFluor-594 Donkey Anti-Goat (Invitrogen lot #1180089, 1:200) or AlexaFluor-594 Goat Anti-Mouse (Invitrogen lot # 1219862, 1:200). Nuclei were counterstained with DAPI. For HOPX over-expression studies (**Figure 5**), cells were fixed in 4%(vol/vol) paraformaldehyde, blocked for an hour with 5% (vol/vol) normal goat serum (NGS), and incubated overnight with primary antibody in 1% NGS, followed by secondary antibody staining in NGS. Measurements of CM area were performed using Image J software. Analysis was done on a Leica TCS-SPE Confocal microscope using a 40x or 63x objective and Leica Software. Primary antibodies used were: αActinin 1:250 Sigma A7811 anti-mouse, Titin 1:300 Myomedix TTN-9 (cTerm) anti-rabbit, GFP 1:300 Invitrogen A-11122 anti-rabbit. Secondary antibodies and other reagents used were: DAPI at a concentration of 0.02μg/mL, phalloidin alexa fluor 568 1:250, alexa fluor 488 or 647-conjugated goat anti-mouse and anti-rabbit secondary antibodies 1:500 (Molecular Probes).

##### Flow Cytometry

Cells were fixed with 4% paraformaldehyde and permeabilized in 0.75% saponin. Cells were labeled for flow cytometry using cardiac troponin T (Pierce, MA5-12960) or APLNR (R&D, FAB856A) and corresponding isotype control. Cells were analyzed using a BD FACSCANTO II (Becton Dickinson, San Jose, CA) with FACSDiva software (BD Biosciences). Data analysis was performed using FlowJo (Tree Star, Ashland, Oregon).

##### Single cell isolation

For each differentiated day 0, 2, 5, 15, and 30 time point, differentiated cells were dissociated with 0.5% EDTA + 0.25% Trypsin (ThermoFisher, 15400054) and neutralized with foetal bovine serum (Scientifix, FFBS-500) and DMEM/F12 media (Life Technologies Australia, 11320033) (1:1 ratio). For each time point, 2 pooled samples were collected, each pool comprised approximately 12 independent differentiation samples. Cells were centrifuged at 1200 rpm for 4 minutes and resuspended in Dulbecco’s PBS (Gibco; 14190) with 0.04% bovine serum albumin (Sigma Aldrich, B6917) and immediately transported for scRNA-Seq processing. Viable cells were sorted using a Propidium Iodide stain and retained on ice in Dulbecco’s PBS + 0.04 % bovine serum albumin. A Countess automated counter (Invitrogen) was used to check final cell viability using Trypan Blue exclusion.

##### Single cell RNA-Sequencing

A Chromium instrument (10X Genomics, Millennium Sciences) was used to partition sorted, viable cell suspensions (8×10^5^-1×10^6^ cells/mL) into single cell droplets using the Single Cell 3′ Library, Gel Bead and Multiplex Kit (version 1; 10X Genomics; PN-120233) as per the manufacturer’s protocol. Each time point was run in duplicate, resulting in 10 sample preparations. The samples were loaded into Single Cell 3′ chips (10X Genomics) at a concentration optimized to capture approximately 5,000 cells in individual 1-cell droplets. Single cell libraries were sequenced using an Illumina NextSeq 500 instrument as previously described (Nguyen et al., in review-b).

##### Bioinformatics processing

Bioinformatics mapping of reads to original transcripts and cells was by *cellranger* pipeline v1.3.1 (*mkfastq, count, aggr*) using GRC38p7 human reference genome using the STAR software (Dobin et al., 2013). Raw sequencing data between two biological replicates in each timepoint were aggregated and normalised by a subsampling procedure (Zheng et al., 2017). Methods for data preprocessing to filter outlier genes and cells based on median absolute deviation, and data dimensionality reduction by PCA and tSNE normalization are previously described in Nguyen et al (Nguyen, 2018) and as is implemented in the R *ascend* package (Senabouth et al., 2018). After sample-to-sample normalisation and data filtering, cell-to-cell normalisation was done by scran (Lun et al., 2017). Clustering algorithm was described in Nguyen et al (2017 and 2018) and is implemented in the ascend as well as the *scGPS* R packages. Furthermore, for reproducibility and broader usability of the valuable data resource, we have submitted all data to Array Express and created a web database resource with interactive data mining tools for users to explore the entire dataset without requirement for programming.

The clustering algorithm, CORE, builds a high-resolution clustering tree structure, dynamically groups branches of the tree at 40 different height cutoffs, and compares clustering results by using adjusted Rand indexes to find a stable clustering point (Nguyen et al., in review-a). The optimal clustering point meets two criteria, including robust to changing parameters and less different from a reference with the highest number of clusters. CORE algorithm is described in details in Nguyen et al (2017) and is implemented in the ascend R package (Senabouth et al., 2018). We validated the clustering results using multiple dimensionality reduction methods (PCA, tSNE, MDS), imputation method (CIDR), and pseudotime mapping (Diffusion and Monocle).

The processed data. post filtering, normalisation, and clustering was used as the input for differential expression analysis. We performed DESeq (Anders and Huber, 2010) differential expression analysis between cells in one subpopulation compared to all remaining cells in a given time-point. We observed that DESeq performed better than DESeq2, consistent to the report by Dal Molin et al(Dal Molin et al., 2017). scran normalised counts were used as input to DESeq with adjustments to add and subtract pseudocount before and after DE analysis to better estimate fold changes. Bonferroni correction as applied to account for multiple testing.

##### Bulk RNA-Sequencing

hiPSC-CMs were harvested for RNA preparation and genome wide RNA-seq (>20 million reads). RNA-seq samples were aligned to hg19 using Tophat, version 2.0.13 (Trapnell et al., 2009). Gene-level read counts were quantified using htseq-count (Anders et al., 2015) using Ensembl GRCh37 gene annotations. Genes with total expression above 1 normalized read count across RNA-seq samples in each binary comparison (e.g., HOPX vs. control) were kept for differential analysis using DESeq (Anders and Huber, 2010). Princomp function from R was used for Principal Component Analysis. TopGO R package (Alexa et al., 2006) was used for Gene Ontology enrichment analysis.

##### Protein extraction and western blot analysis

Cells were lysed directly on the plate with a lysis buffer containing 20mM Tris-HCl pH 7.5, 150mM NaCl, 15% Glycerol, 1% Triton X-100, 1M β-Glycerolphosphate, 0.5M NaF, 0.1M Sodium Pyrophosphate, Orthovanadate, PMSF and 2% SDS (Moody et al., 2017). 25U of Benzonase Nuclease (EMD Chemicals, Gibbstown, NJ) was added to the lysis buffer right before use. Proteins were quantified by Bradford assay (Bio-rad), using BSA (Bovine Serum Albumin) as Standard using the EnWallac Vision. The protein samples were combined with the 4x Laemmli sample buffer, heated (95°C, 5min), and run on SDS-PAGE (protean TGX pre-casted 4%-20% gradient gel, Bio-rad) and transferred to the Nitro-Cellulose membrane (Bio-Rad) by semi-dry transfer (Bio-Rad). Membranes were blocked for 1hr with 5% milk and incubated in the primary antibodies overnight at 4°C. The membranes were then incubated with secondary antibodies (1:10000, goat anti-rabbit or goat anti-mouse IgG HRP conjugate (Bio-Rad) for 1hr and the detection was performed using the immobilon-luminol reagent assay (EMD Millipore). Primary antibodies are as follows: Alpha tubulin antibody Cell Signalling Technologies (2144) 1:2000 and anti-GFP Invitrogen (A-11122) anti-rabbit 1:1000.

##### Genomics data sets

Previously published ChIP-seq and gene expression data sets were analyzed for this study. Analysis of cardiac differentiation chromatin dynamics and gene expression by RNA-seq were published previously (Kuppusamy et al., 2015; Palpant et al., 2017b) with data accessed from GEO GSE97080. HOPX gene expression analysis were derived from Stemformatics (Wells et al., 2013) using the following data sets: a dual reporter MESP1-mCherry/NKX2-5 GFP reporter hESC line at day 0 and day 3 of directed differentiation sorted for MESP1 positive vs. negative cells (Stemformatics ID: Hartogh_2015_25187301) (Den Hartogh et al., 2015) and human foetal heart samples isolated at each of three trimesters comparing ventricle and atrial expression (Stemformatics ID:
van_den_berg_2015_26209647) (van den Berg et al., 2015). Human fetal heart gene expression data were downloaded from ENCODE (experiment #: ENCSR047LLJ, ENCSR863BUL, ENCSR769LNJ, ENCSR433XCV, and ENCSR675YAS). HOPX expression in engineered tissue, adult heart tissue, and hPSC-CMs were acquired from previous work by Mills et al (Mills et al., 2017).

##### Gene ontology visualization

Gene ontology analysis was performed using DAVID with significance threshold set at FDR < 0.05. The p-values from gene ontology analysis were visualized using the R package corrplot (Wei and Simko, 2016), where the radius of the circle is proportional to the negative natural log of the input p-value.

##### Spearman correlation analysis

We obtained FACS sorted bulk cardiac subtypes (Quaife-Ryan et al., 2017) and single-cell RNA-seq data generated from developing mouse heart (Li et al., 2016). The normalized expression data from these two sources was merged with our scRNA-seq expression data. Mouse Ensembl IDs were converted to human ortholog gene IDs and a new expression matrix was generated using only the 13,490 genes common to all three datasets. Spearman’s rank correlation was used to compare the expression levels of genes between samples, and the significance of the differences between pairs of correlation coefficients were calculated using a Fisher Z-transformation.

##### iTranscriptome sample preparation and data analysis

Samples were generated according to the methodology published in (Peng et al., 2016). E6.5, E7.0 and E7.5 embryos were cryo-sectioned along the proximal-distal axis. Populations of approximately 20 cells were collected from different regions of the cross-section by laser microdissection and processed for RNA sequencing. Two sets of embryos for each embryonic age were dissected (**Figure S1**): the first set from the epiblast - E6.5: anterior and posterior; E7.0: anterior, left, right and posterior; E7.5: anterior, left anterior and posterior, right anterior and posterior, posterior. The second set from the three germ layers - E6,5: posterior epiblast and endoderm: anterior and posterior; E7.0: posterior epiblast, mesoderm: anterior and posterior, and endoderm: anterior and posterior; E7.5: posterior epiblast, mesoderm: anterior and posterior, and endoderm: anterior and posterior. Differentially expressed genes (DEGs) were screened first by unsupervised hierarchical clustering method to group samples in the respective regions. Genes with an expression level FPKM>1 and a variance in transcript level across all samples greater than 0.05 were selected. To identify inter-region specific DEGs, each of these selected genes was submitted to a t-test against the level of expression in the other regions. Genes with a p.value< 0.01 and a fold change >2.0 or <0.5 were defined as DEGs. The gene expression pattern (region and level of expression by transcript reads) of the gene of interest was mapped on the corn plots, where each kernel represents the cell population sampled at a defined position in the germ layers, to generate a digital rendition of whole mount in situ hybridization.

##### Constructing regulatory differentiation networks scdiff

Detailed computational model and derivation for *scdiff* are provided in (Ding et al., In review). *scdiff* software is available on GitHub (https://github.com/phoenixding/*scdiff*;). The method is initialized using Spectral clustering based on cell-to-cell Spearman correlation, followed by an ensemble strategy to determine the optimal K clusters. The model then iteratively connects clusters (representing states in the probabilistic model) between time points using a “Similarity To Ancestor-STA” strategy (with day 0 as the first time-point), based on expression similarity (Spearman correlation). Cell reassignment is based on a Kalman filter probabilistic model. The initial set of states and their connectivity is iteratively updated by learning trajectory and branching models constrained by transcription factor (TF)-gene interactions via a logistic regression classifier to maximize the ability to predict the expression of target gene based on the interaction data. Details of the implementation of *scdiff* are outlined below.

##### Initial clustering of single cells

*scdiff* starts by clustering the cells in each of the time points measured. While the original *scdiff* method used spectral clustering, this method was unable to scale for the large number of cells profiled in this study. We have thus revised scdiff for this study by changing the original clustering methods to a more efficient method. Specifically, we used PCA with 10 dimensions followed by K-means for the initial clustering which led to faster runtime while not greatly affecting performance. To determine the initial number of clusters (k) for each time point we combine 3 widely used clustering quality assessment scores: the Silhouette Score (Rousseeuw, 1987), Davis-Bouldin index (Davies and Bouldin, 1979) and AIC (Akaike, 1998) (Akaike information criterion). We used a bootstrapping strategy to combine these. We first selected a random subset of 90% of the genes. Next, we calculate the Silhouette score, Davis Bouldin score and AIC scores for different k values between 2 and 20 for each time point. We compute a combined score for each of these k values based on the subset of genes selected and repeat this process 100 times (each time with a new random gene subset). We select the optimal k by summing up the scores for each of the possible k values across the 100 repeats.

##### Initial model construction

Initial clustering is based on the time point associated with each cell. However, several recent studies indicate that cells may be unsynchronized with respect to their state even if they are collected at the same time point (Trapnell et al., 2014). Thus, some of the clusters at a specific time point may represent states that are either earlier or later than other clusters in the same time. To address this we next use a correlation-based method to reassign clusters to time points. Once we determined the set of clusters associated with each level (time point), we connect clusters in each level to the most similar cluster (in terms of correlation) at the level directly above it (its parent cluster). This leads to a directed graph with potentially multiple roots (initial set of clusters for the first time points) which structurally represents the initial differentiation model.

##### Predicting TFs regulating differentiation pathways

An important aspect of *scdiff* is the ability to both reconstruct and analyze the differentiation pathways based on the set of TFs that regulate various state transitions. We used bulk TF-gene interaction data from (Ernst et al., 2007; Schulz et al., 2012) for this analysis. Following the initial model construction, we first identify a set of differentially expressed (DE) genes for each cluster (state) in our model. Using this set, we identify TFs that are enriched for DE targets based on the hyper-geometric distribution. Next, we check which of the candidate TFs is expressed in the parent node of the state. TFs that are both significantly enriched and expressed are used in the to define a logistic regression function which modifies the likelihood of assigning cells to the different clusters in the model Thus, cell assignment is based on both, expression similarity to other cells in the state and the expression of targets of TFs predicted to regulate this state.

##### Iterative assignment of cells to states

Given initial assignments of cells and TFs to states, we can compute the MLE of the transition and emission noise variance. We next iterate between two steps. The first uses the parameters learned to reassign cells and TFs to states and the second uses the assigned cells and TFs to re-learn model parameters. During the iterative process some states may become empty and if this happens they are removed from the model. The process stops when it converges (no more cells are reassigned) and the resulting model is returned.

*scdiff* was run with the following parameters: -k auto -l 1 -s 1 -d 1. The expression data matrix was normalized at two levels: by batches (using the 10x cellRanger R package), and then by cells (using the deconvolution method in the ‘SCRAN’ package) in R. A thorough quality check to remove outlier genes and cells (outside 3 × median absolute deviation range) based on mitochondrial, ribosomal genes, library sizes, and number of detected cells (as described in our ASCEND R package - under review).

## Supplemental Figures

**Figure S1 related to Figure 2.**
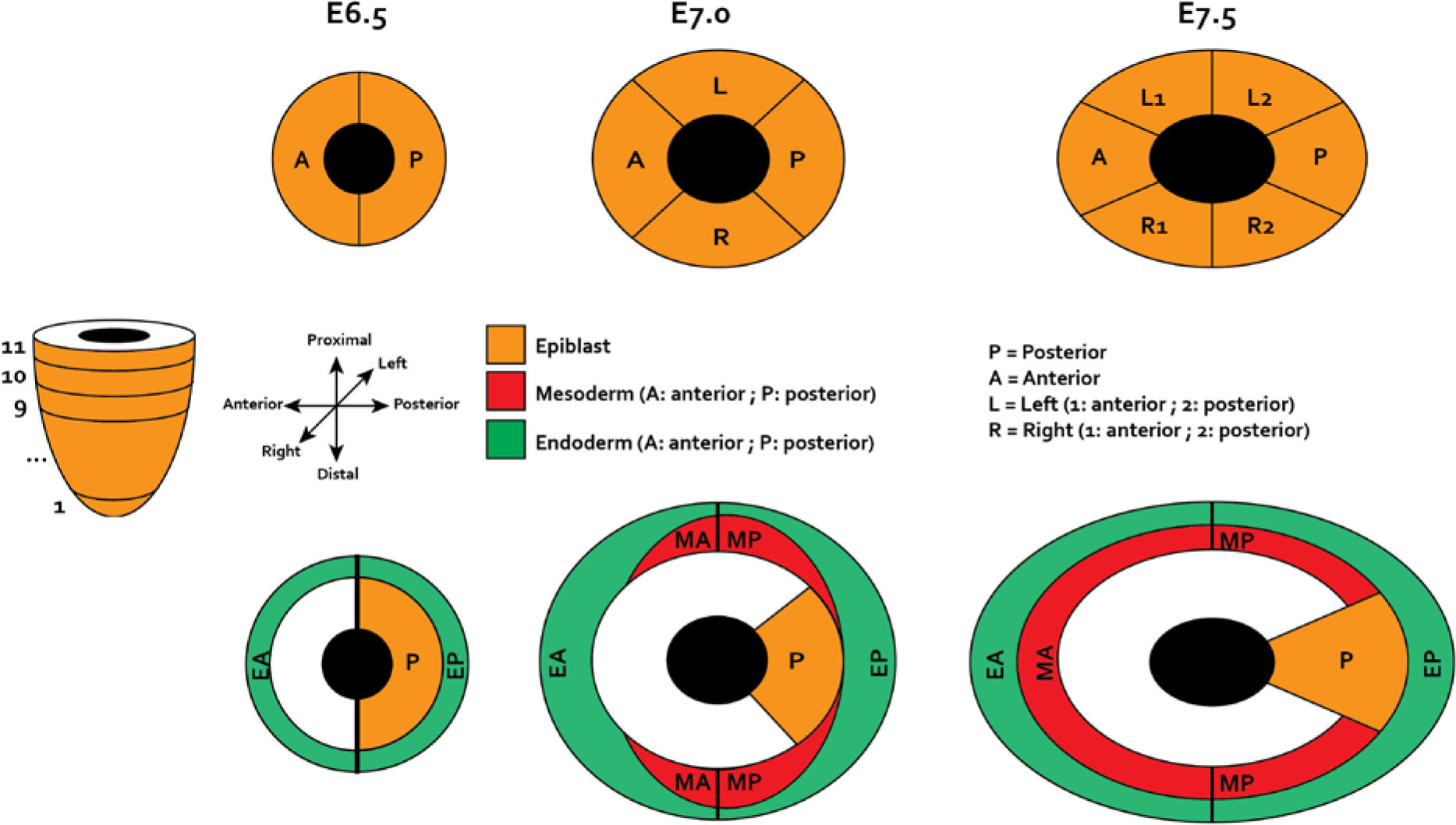
iTranscriptome analysis of the gastrulating mouse embryo. Schematic of the germ layer regions in E6.5, E7.0 and E7.5 embryos from which cell populations were sampled by laser capture microdissection and analyzed by RNA-seq. Sections numbered in distal-proximal order: E6.5: 1 to 7; E7.0: 1 to 11; E7.5: 1 to 9. E6.5, epiblast: anterior (A), posterior (P), Endoderm: anterior (EA), posterior (EP); E7.0, Epiblast: anterior (A), left (L), right (R), posterior (P), Mesoderm: anterior (MA), posterior (MP), Endoderm: anterior (EA), posterior (EP); E7.5: Epiblast: anterior (A), left anterior (L1), left posterior (L2), right anterior (R1), right posterior (R2), Mesoderm: anterior (MA), posterior (MP), Endoderm: anterior (EA), posterior (EP). Triple double-headed arrows indicate the embryonic axes of the gastrula-stage embryo.

**Figure S2 related to Figure 2.**
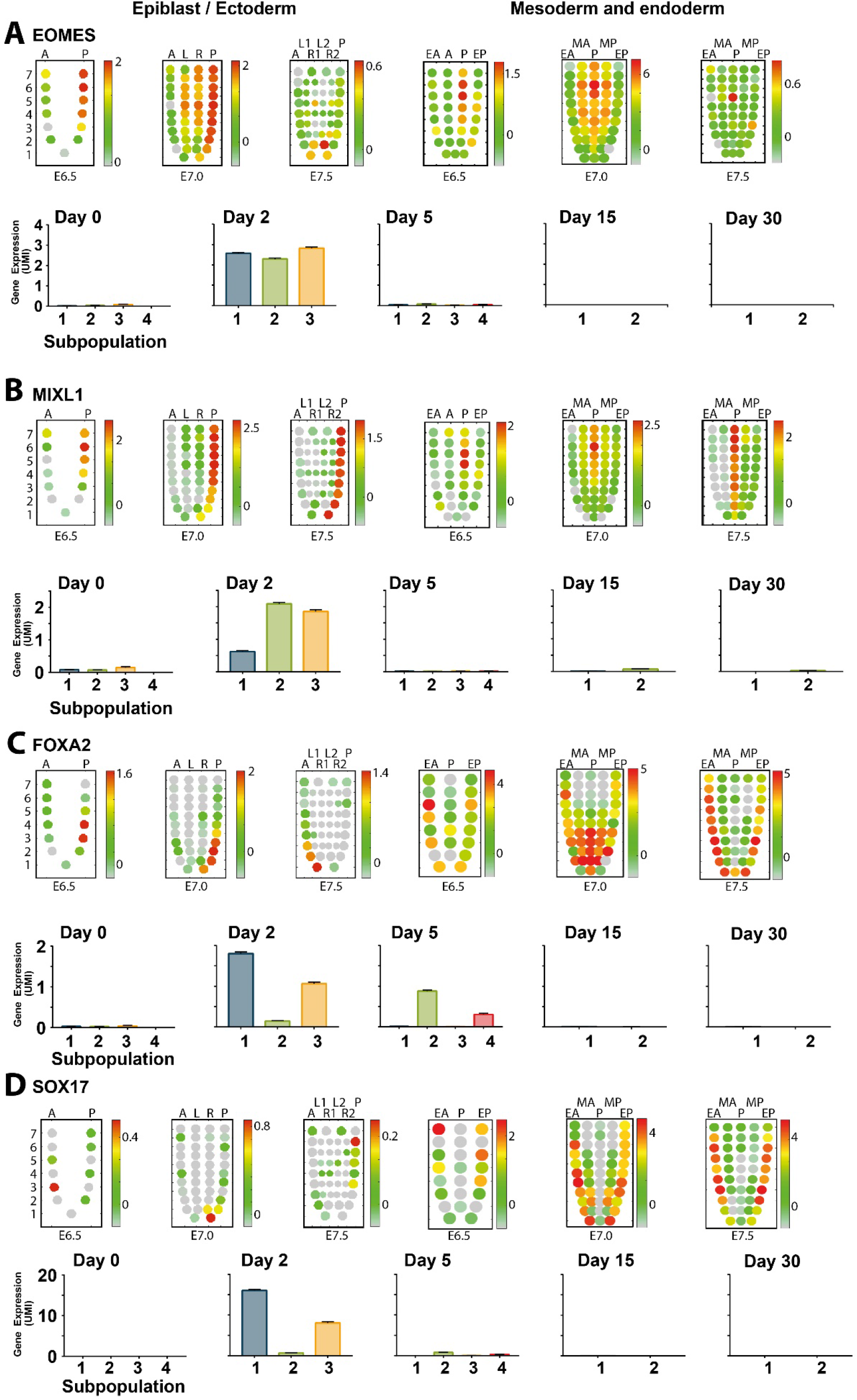
Comparative analysis of single cell expression patterns against spatio-temporal RNA-seq analysis of gene expression in mouse gastrula stage embryos. Corn plots showing the spatial domains of **(A)** EOMES, **(B)** MIXL1, **(C)** FOXA2, and **(D)** SOX17 expression in the epiblast/ectoderm, mesoderm, and endoderm of E6.5, E7.0, and E7.5 mouse embryos during gastrulation (unpublished RNA-seq data for E6.5 (n=6) and E7.5 (n=6) embryos and published data for E7.0 embryo, (Peng et al., 2016)). Positions of the cell populations (so-called “kernels” in the 2D plot of RNA-Seq data) in the germ layers: the proximal-distal location in descending numerical order (1 = most distal site); in the transverse plane of the epiblast/ectoderm - E6.5, Anterior (A) and Posterior (P) quadrants; E7.0, Anterior (A), Posterior (P), Left (L), and Right (R) quadrants; E7.5, Anterior (A) and Posterior (P) quadrants, Left-anterior half (L1), Right-anterior half (R1), Left-posterior half (L2), and Right-posterior half (R2) of left and right quadrants; in the transverse plane of the mesoderm and endoderm - Anterior half (EA) and Posterior half (EP) of the endoderm, Anterior half (MA) and Posterior half (MP) of the mesoderm, and Posterior epiblast (P) containing the primitive streak. Color scales represent levels of expression as log10 of fragments per kilobase million (FPKM+1). Below each set of corn plots, the mean expression value of each gene during cardiac directed differentiation in each subpopulation over the time course of lineage decisions are shown. UMI: Unique Molecular Identifier.

**Figure S3 related to Figure 2.**
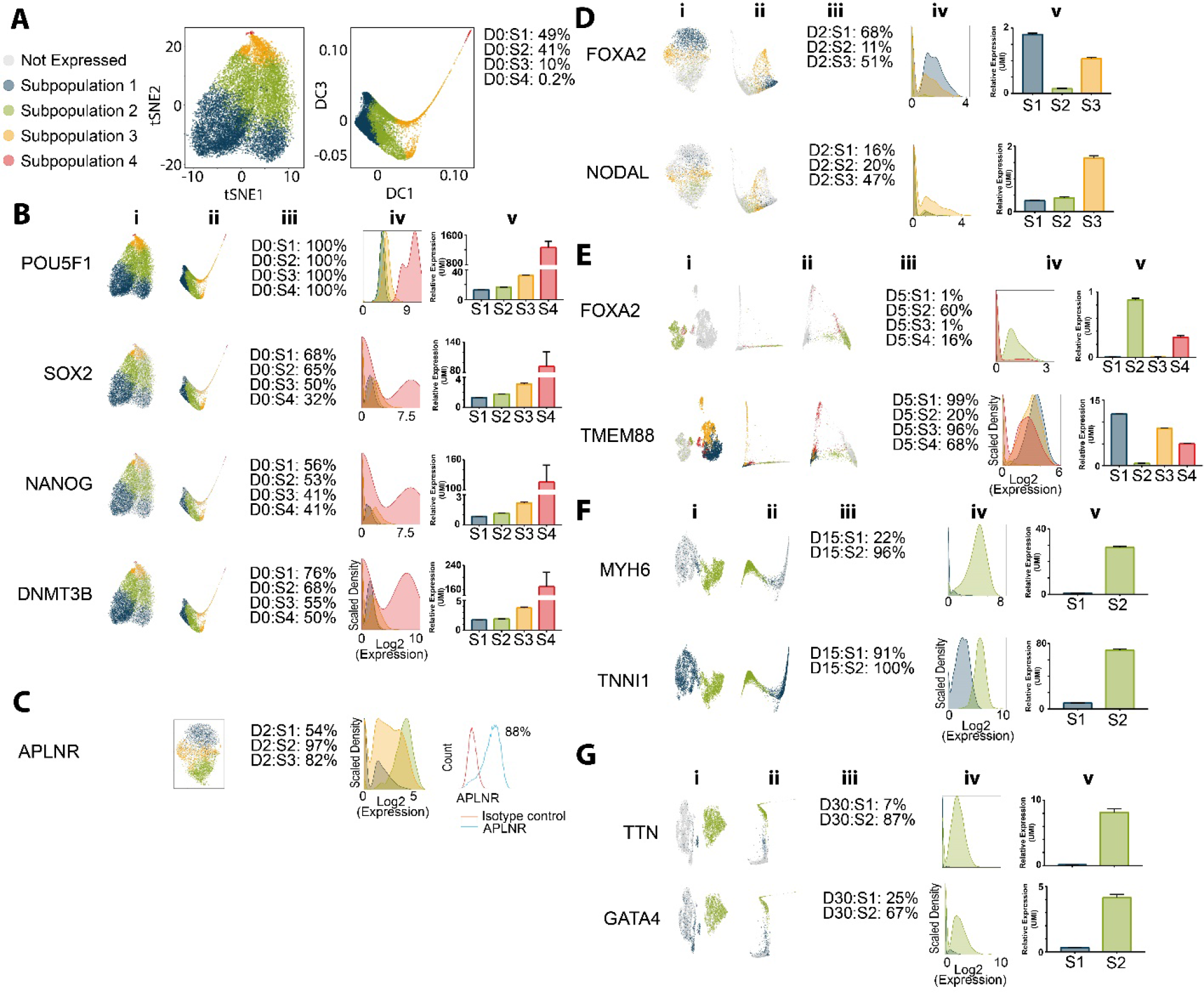
Analysis of lineage-specific genes identified at specific time points of differentiation to establish subpopulation identities. **(A-B)** Analysis of day 0 subpopulations represented by *t*-SNE (left) and diffusion (right) plots and the percent of cells in each subpopulations (D0:S1-S4), **(B)** analysis of pluripotency genes POU5F1, SOX2, NANOG, and DNMT3B (data from left for each gene: **(i)** *t*-SNE plot and **(ii)** diffusion plot showing cells +/− for gene, **(iii)** percent of cells expressing gene, **(iv)** histogram of gene expression, and **(v)** expression level of gene in each subpopulation). **(C)** Analysis of computational analysis of the mesoderm marker APLNR on day 2 of differentiation from single cell RNA-seq data (left) compared to FACS analysis of APLNR (right). **(D-G)** Analysis of expression patterns and dynamics of genes associated with **(D)** mes-endoderm specification at day 2 of differentiation (definitive endoderm transcription factor FOXA2 and anterior primitive streak mes-endoderm transcription factor NODAL), **(E)** the progenitor stage of differentiation at day 5 (definitive endoderm transcription factor FOXA2 and cardiac progenitor Wnt regulatory gene TMEM88), early cardiomyocyte stage of differentiation at day 15 (fetal sarcomere isoform of myosin MYH6 and troponin I TNNI1), and **(G)** late stage of cardiomyocyte differentiation at day 30 (sarcomere gene TTN and cardiac transcription factor GATA4) (data from left for each gene: **(i)** t-SNE plot and **(ii)** diffusion plot showing cells +/− for gene, **(iii)** percent of cells expressing gene, **(iv)** histogram of gene expression, and **(v)** expression level of gene in each subpopulation). UMI: Unique Molecular Identifier.

**Figure S4 related to Figures 3 and 4.**
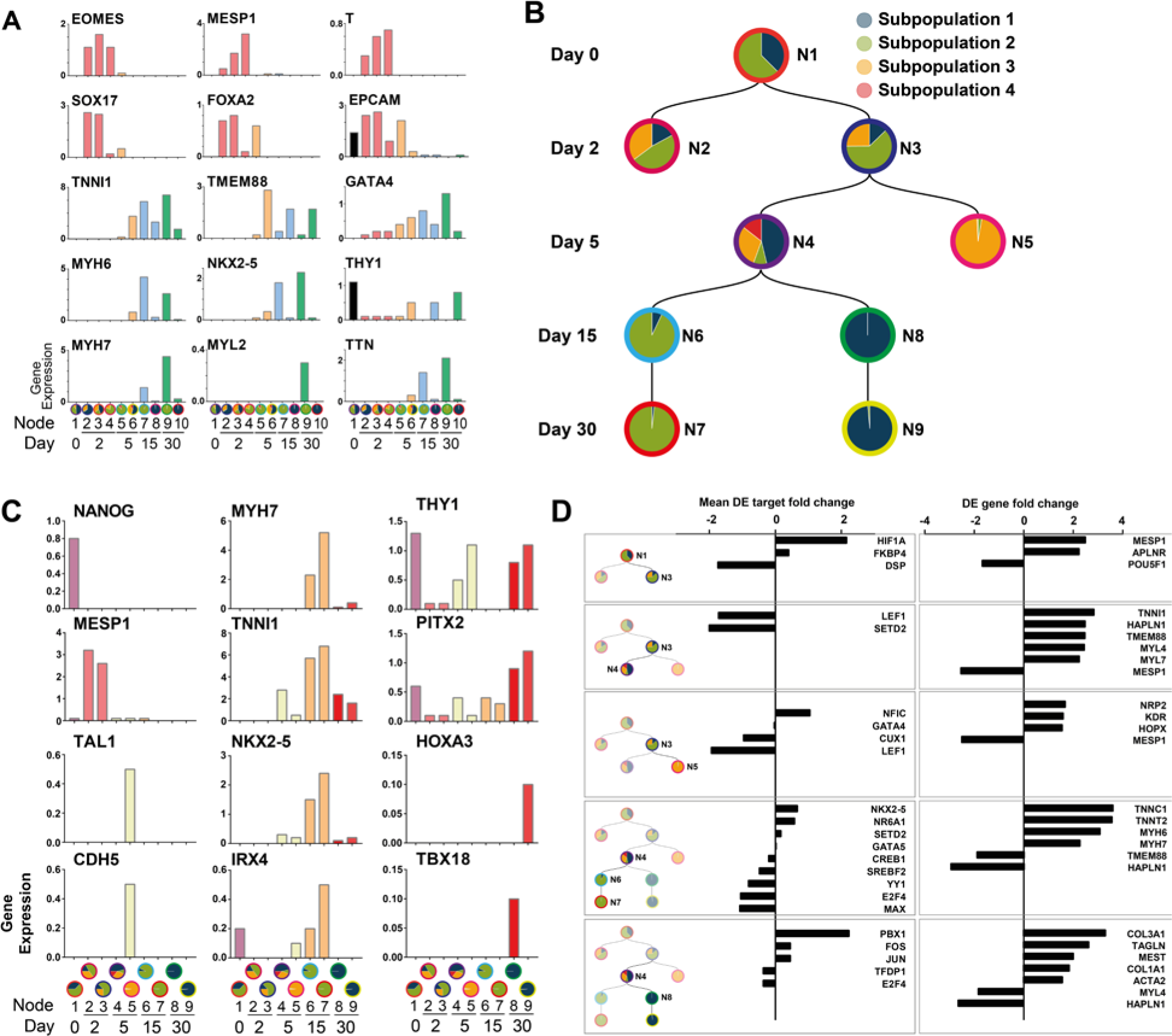
Lineage prediction analysis of cardiac differentiation by scdiff. **(A)** Expression level within each node for known cardiac and endodermal lineage markers reflecting *in vivo* populations. Gene expression is represented across all nodes: EOMES (pan-mesoderm), MESP1 and T (cardiogenic mesoderm), SOX17, FOXA2, EPCAM (endoderm), TMEM88, GATA4, TNNI1 (cardiac progenitors), THY1 (fibroblasts), MYH6, MYH7, MYL2, TTN, NKX2-5 (late-stage cardiac differentiation markers). **(B)** Lineage tracing prediction of only HOPX cells at each stepwise transition from pluripotency into cardiac lineages predicted on the basis of gene regulatory networks (GRN) detected between pairwise changes in cell state during differentiation. Circles indicate distinct nodes governed by a common GRN. Since cells can be re-assigned based on network stability, the redistribution of subpopulations established by our initial unsupervised clustering analysis (as outlined in **Figure 2**) are represented as pie charts within each circle indicating the percent of cells from each subpopulation contributing to that node. Nodes are demarcated as N1-N9. **(C)** Expression level within each node for known developmental lineage markers. Gene expression is represented across all nodes: NANOG (pluripotency), MESP1 (cardiac-mesoderm), TAL1 and CDH5 (endothelium) TNNI1, MYH7, NKX2-5, IRX4 (primarily first heart field cardiac genes), THY1, PITX2, HOXA2, and TBX18 (second heart field and outflow tract development). **(D)** Expression level within each node for known pluripotency, mes-endodermal, and cardiac genes in HOPX cells comprising different stages of differentiation as outlined in Figure 4F-G and **Table S5**. The analysis identifies transcription factors and genes controlling stage-specific regulatory networks underlying cell fate transitions. Mean DE target fold change calculates the fold change for the differentially expressed targets of the TF. DE gene fold change shows up or down-regulated fold change of TF target genes.

**Figure S5 related Figure 4.**
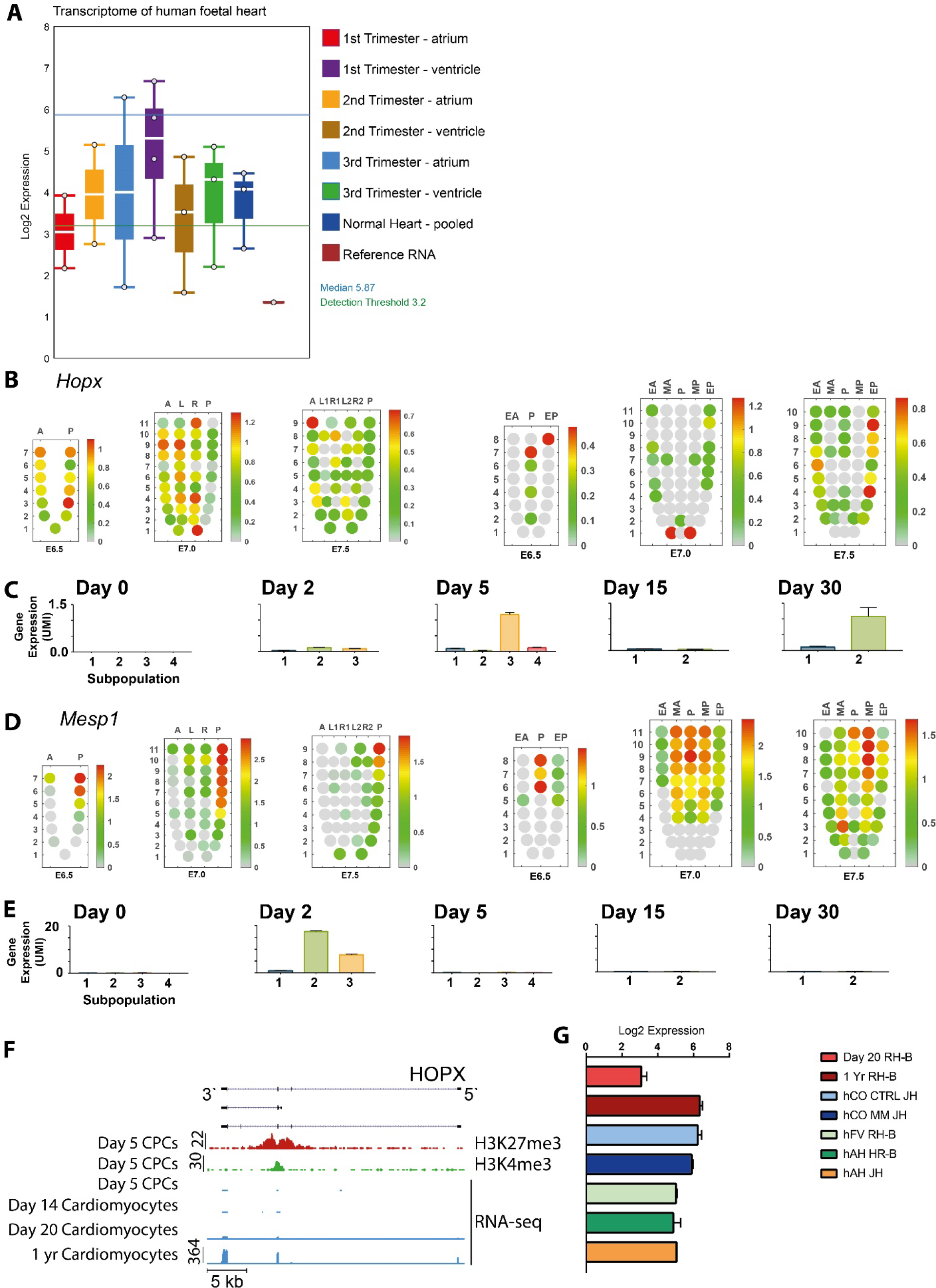
Analysis of HOPX expression during cardiac development *in vitro* and *in vivo*. **(A)** Expression of HOPX from human foetal heart samples isolated at each of three trimesters comparing ventricle and atrial expression. **(B-E)** Corn plots showing the spatial domains of **(C)** HOPX and **(E)** MESP1 expression in the epiblast/ectoderm, mesoderm, and endoderm of E6.5, E7.0, and E7.5 mouse embryos during gastrulation (unpublished RNA-seq data for E6.5 (n=6) and E7.5 (n=6) embryos and published data for E7.0 embryo, (Peng et al., 2016)). Positions of the cell populations (so-called “kernels” in the 2D plot of RNA-Seq data) in the germ layers: the proximal-distal location in descending numerical order (1 = most distal site); in the transverse plane of the epiblast/ectoderm - E6.5, Anterior (A) and Posterior (P) quadrants; E7.0, Anterior (A), Posterior (P), Left (L), and Right (R) quadrants; E7.5, Anterior (A) and Posterior (P) quadrants, Left-anterior half (L1), Right-anterior half (R1), Left-posterior half (L2), and Right-posterior half (R2) of left and right quadrants; in the transverse plane of the mesoderm and endoderm - Anterior half (EA) and Posterior half (EP) of the endoderm, Anterior half (MA) and Posterior half (MP) of the mesoderm, and Posterior epiblast (P) containing the primitive streak. Color scales represent levels of expression as log10 of fragments per kilobase million (FPKM+1). Below each set of corn plots are shown the mean expression value of HOPX **(B-C)** and MESP1 **(D-E)** during cardiac directed differentiation in each subpopulation over the time course of lineage decisions. **(F)** ChIP-seq and RNA-seq analysis of the HOPX locus in day 5 cardiac progenitor cells showing chromatin levels for H3K4me3 vs H3K27me3 and gene expression by RNA-seq of cardiomyocyte differentiation from day 5 to 1 year old differentiated cardiomyocytes. **(G)** Human hPSC-monolayer derived cardiomyocytes at 20 days (day 20 RH-B, n=3) and 1 year of differentiation (1 yr RH-B, n=3), h cardiac organoids in control (hCO CTRL JH, n=4) and maturation media (hCO MM JH, n=4), human fetal ventricle (hFV RH-B, n=2), and human adult heart (hAH HR-B, n=2 and hAH JH, n=1). UMI: Unique Molecular Identifier.

**Figure S6 related to Figure 4.**
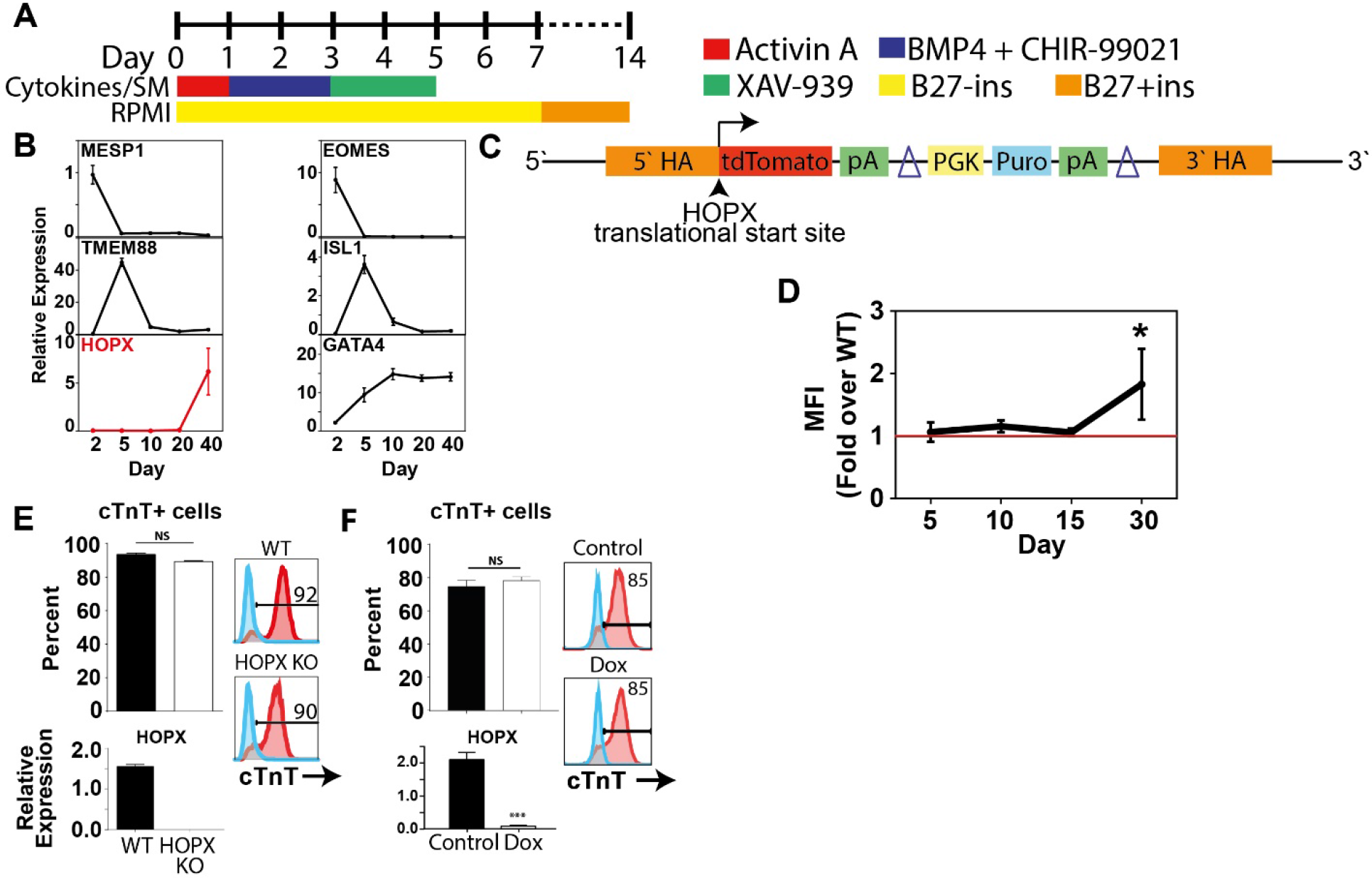
HOPX expression in Activin A/BMP4 mediated directed differentiation from pluripotency. **(A)** Schematic of protocol for Activin A/BMP4 differentiation (ABCX-100) from pluripotency as described previously (Palpant et al., 2017a). **(B)** Gene expression profiling by qRT-PCR during Activin A/BMP4 differentiation using hESC line RUES2 showing proper temporal dynamics of expression for mesoderm at day 2 (MESP1, EOMES) cardiac progenitor at day 5 (TMEM88, ISL1) and GATA4 activated between days 5-40 compared to the temporal expression of HOPX. **(C)** Schematic of the HOPX-tdTomato reporter cell line in which the fluorescence reporter is knocked into the TSS of the HOPX locus for real-time dynamic analysis of HOPX activity during differentiation (Palpant et al., 2017b). **(D)** Mean fluorescence intensity analysis of HOPX reporter cells (normalized to WT controls) during a time course of cardiac directed differentiation using Activin A/BMP4 induction. **(E-F)** FACS analysis of HOPX loss of function stem cell lines including HOPX genetic KO **(E)** and HOPX CRISPRi **(F)** showing no difference in cardiac specification in HOPX deficient cells.

**Figure S7 related to Figure 5.**
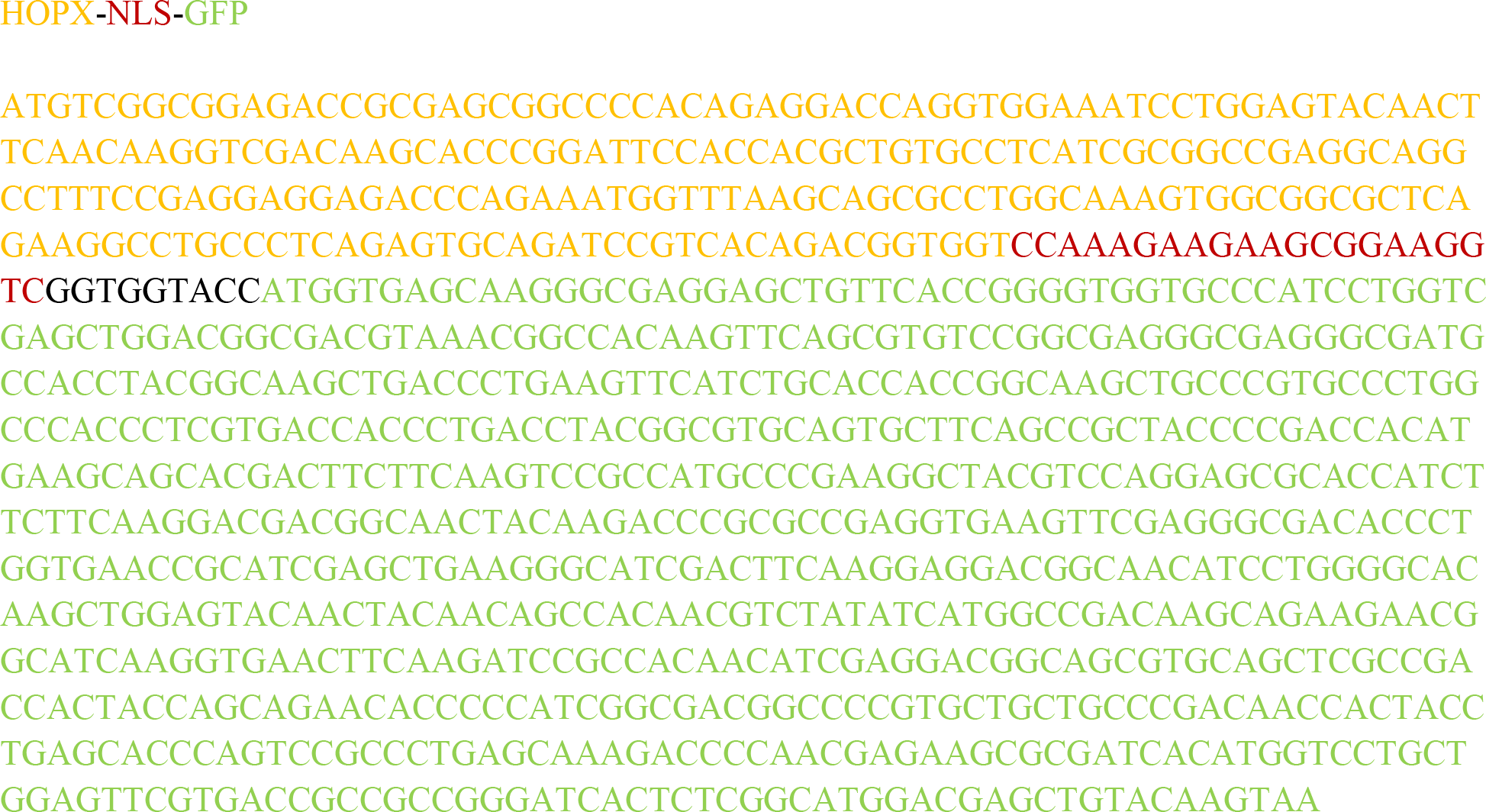
HOPX-NLS-eGFP construct sequence.

## Supplemental Tables

**Table S1.** Differential gene expression and gene ontology analysis of subpopulations identified during cardiac directed differentiation at single cell resolution.

**Table S2**. Computational analysis of transcription factor and gene regulatory networks underlying *scdiff* prediction of lineage trajectories for the full single cell time course data set.

**Table S3.** Transcription factor and epigenetic regulators of cardiovascular fate diversification analyzed for expression in eleven subpopulations comprising cell fate transitions from day 2, 5, 15, and 30.

**Table S4.** Analysis of HOPX expression in various cell types from *in vivo* heart development.

**Table S5.** Computational analysis of transcription factor and gene regulatory networks underlying *scdiff* prediction of lineage trajectories for only HOPX^+^ cells during the full time course data set.

**Table S6.** RNA-seq analysis of HOPX over-expression in cardiomyocytes.

**Table S7.** Gene ontology analysis of differentially expressed genes in HOPX over-expression vs control. Table S8. Statistical analysis of gene expression shown in heat maps in **Figure 6J-M.**

**Table S9.** Primer sequences for HOPX CRISPRi and qRT-PCR analysis.

